# Benchmarking long-read sequencing strategies for obtaining ASV-resolved rRNA operons from environmental microeukaryotes

**DOI:** 10.1101/2023.09.21.558928

**Authors:** Christina Karmisholt Overgaard, Mahwash Jamy, Simona Radutoiu, Fabien Burki, Morten Kam Dahl Dueholm

## Abstract

The use of short-read metabarcoding for classifying microeukaryotes is challenged by the lack of comprehensive 18S rRNA reference databases. While recent advances in high-throughput long-read sequencing provide the potential to greatly increase the phylogenetic coverage of these databases, the performance of different sequencing technologies and subsequent bioinformatics processing remain to be evaluated, primarily because of the absence of well-defined eukaryotic mock communities. To address this challenge, we created a eukaryotic rRNA operons clone-library and turned it into a precisely defined synthetic eukaryotic mock community. This mock community was then used to evaluate the performance of three long-read sequencing techniques (PacBio HiFi, and Nanopore UMI with/without clonal pre-amplification) and three tools for resolving amplicons sequence variants (ASVs) (Uchime3, Unoise3, and DADA2). We investigated the sensitivity of the sequencing techniques based on the number of detected mock taxa, and the accuracy of the different ASV-calling tools with a specific focus on the presence of chimera among the final rRNA operon ASVs. Based on our findings, we provide recommendations and best practice protocols for how to cost-effectively obtain essential error-free rRNA operons in high-throughput. An agricultural soil sample was used to demonstrate that the sequencing and bioinformatic results from the mock community also translates to highly diverse natural samples.

## Introduction

Community composition studies of microeukaryotes are often based on short-read metabarcoding where the V4 or V9 regions of the 18S rRNA genes are commonly used for protists (Singer et al., 2021; Vaulot et al., 2022), whereas the ITS1 or ITS2 regions are applied for fungi (Tedersoo et al., 2014; Tkacz et al., 2020). Short-read metabarcoding has revolutionized the way we investigate microbial communities. Nonetheless, such short reads come with constraints. Their capacity for both taxonomic and phylogenetic discernment is limited, making them reliant on reference databases populated with full-length or near full-length 18S rRNA genes. The recent years’ advancement in long-read sequencing and bioinformatic pipelines creating high-quality sequences (Callahan et al., 2019; Jamy et al., 2020; Karst et al., 2021) has facilitated the generation of millions of full-length 16S rRNA reference sequences (Dueholm et al., 2022; Dueholm et al., 2020) and even bacterial rRNA operons (Martijn et al., 2019). The high-quality reference sequences can be used to improve classification and determine the primer bias associated with commonly used primers for metabarcoding. However, only a few studies have leveraged these advancements to sequence the rRNA operon of microeukaryotes (Heeger et al., 2018; Jamy et al., 2020, 2022; Latz et al., 2022).

Several long-read sequencing techniques have recently been developed such as PacBio and Nanopore sequencing. PacBio’s HiFi sequencing has become popular because of its ability to produce highly accurate long-read data. The method uses circular consensus sequencing (CCS) to sequence a circular DNA molecule multiple times, producing a precise consensus sequence. Because of the high accuracy and long read length the method has been widely used within fields of human gut research, environmental research, and medical studies (Jamy et al., 2020; Kim et al., 2022; Veiga et al., 2022). Oxford Nanopore has gained immense popularity due to its affordability and easy accessibility, making it another favored technique. Nanopore sequencing utilizes a nanopore-based technology to directly read the DNA molecules as they pass through membrane-embedded pores. This technology allows for real-time sequencing, providing fast and accurate results. It has been used in various fields, including medical research, environmental, and agricultural studies (Michaelsen et al., 2022; Parker et al., 2017; Seitz et al., 2021).

An approach for obtaining highly accurate sequences from Nanopore data was recently introduced (Karst et al., 2021). However, this method requires the use of unique molecular identifiers (UMIs) to correct for the higher error rate in raw Nanopore data. UMIs are short random sequences of nucleotides that are added to DNA or RNA molecules as unique barcodes prior to amplification and sequencing (Figure S1). The purpose of UMIs is to distinguish between PCR duplicates, which are identical DNA fragments generated during the amplification process, and unique molecules in the original sample. After sequencing, reads that have the same UMI are considered PCR duplicates and are collapsed into a single consensus sequence. Addition of UMIs to both ends of the original molecule furthermore enables direct chimera detection as sequences with non-matching UMI pairs are detected as chimeras. UMIs are typically incorporated into a primer used for tagging target molecules and are then retained through the entire sequencing process.

All sequencing methodologies inevitably produce sequences with some degree of error. It’s crucial to differentiate these errors from genuine biological variants. This differentiation is achieved using bioinformatic tools, specifically amplicon sequence variant (ASV) resolving tools. Historically, these tools were designed for short-read data. This original intent might influence their effectiveness when handling long-read data, which has distinct error rates and types. Yet, with the enhanced accuracy brought about by CCS and UMIs, these tools may become suitable for both PacBio and Nanopore data. In fact, tailored pipelines like DADA2 for PacBio CCS data have been developed (Callahan et al., 2019). However, there is no existing DADA2 pipeline for Nanopore data, and concerns have been raised regarding the use of error models for inferring sample composition for error-prone data.

With the growing use of long-read sequencing, it is crucial to assess various sequencing methods and bioinformatic pipelines for generating ASVs to ensure the final sequences represent true biological variants, especially if they are to be used for reference databases. However, such evaluations have not been possible to make for microeukaryote rRNA operons due to the lack of publicly available eukaryotic mock communities.

Here we created a highly diverse synthetic eukaryotic mock sample based on a plasmid-based clone library of full-length rRNA operons. The mock community was used to compare the efficacy of PacBio and Nanopore in recovering mock taxa, evaluate the potential of DADA2, Unoise3, and Uchime3 in producing error- and chimera-free rRNA operon ASVs, and identify the optimal tools and settings for maximizing the number of ASVs obtained. Finally, we determined the applicability of sequencing and bioinformatic results from the mock community to highly diverse natural samples.

## Materials and methods

### General molecular methods

DNA concentration, purity, and quality was measured using the Qubit™ dsDNA HS assay kit, a Thermo Scientific NanoDrop One, and an Agilent 2200 Tapestation with D5000 screentapes (Agilent), respectively, unless otherwise stated. DNA cleanup was performed using CleanNGS beads (CleanNA) in accordance with the manufacturer’s protocol using a bead to sample ratio of 0.7x and elution in 20µL nuclease-free water, unless otherwise stated. All primer and oligos used can be found in Table 1.

**Table 1:**
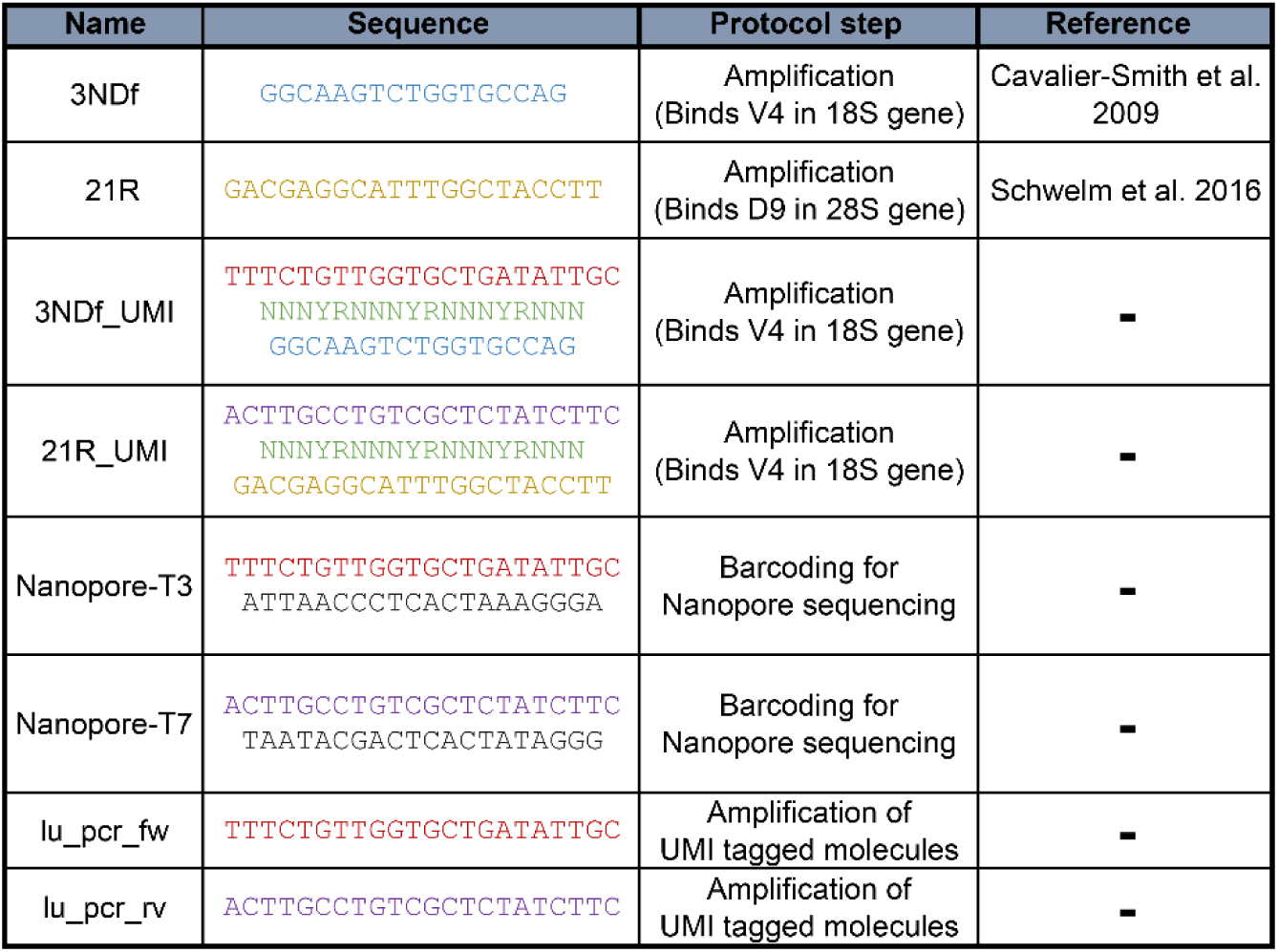
Oligos used in this study. Colors indicate identical sequences.

### Soil and DNA extraction

The Askov soil sample was obtained from the Askov Experimental Station situated in Southern Denmark (GPS coordinates: 55.470 N, 9.111 E) and represents a mix of soils from field B3, plot 315 (no fertilizer), plot 333 (1 NPK), plot 354 (1/2 NPK), and plot 362 (1 1/2 NPK). These fields are part of the Askov Long-Term Experiment established in 1894. The soil is a light sandy loam with 11% clay, 13% silt and 76% sand and can be classified as Alfisol (Typic Hapludalf, USDA Soil Taxonomy). The soil grows a four-course rotation of winter wheat, silage maize, spring barley and a grass-clover mixture used for cutting. Addition of lime every 4-5 years keeps soil pH in the range 5.5 to 6.5 (Christensen et al., 2019). Samples were taken January 4^th^, 2022, from 0-20 cm depth and received the next day. Two teaspoons of soil from each sample were mixed until a homogeneous soil color and texture was obtained. An additional mixing was performed using a spoon in a sterilized container.

DNA was extracted using DNeasy PowerSoil Pro Kit (Qiagen) according to the manufacturer’s protocol and stored at −80°C. DNA cleanup was performed, as previously described, using a bead to sample ratio of 1:1 and eluted in the same amount of nuclease-free water as input. This was done to remove any remaining salt from the DNA extraction. Concentration, purity, and quality was checked as described above. Quality was checked on a genomic screentape (Agilent). DNA extracted from this soil sample was used both for the mock community and as the natural soil sample.

### Preparation of a synthetic eukaryotic mock community

#### PCR amplification of rRNA operons

Eukaryotic rRNA operons (∼4500 bp) from Askov soil was PCR amplified using 20 ng of input DNA, 10 µL of 5X SuperFi buffer, 1 µL of 10 mM dNTP mix, 2.5 µL of 10 µM 3NDf primer, 2.5 µL of 10 µM 21R primer, 10 µL SuperFi GC enhancer, 0.5 µL of 2 U/µL Platinum SuperFi DNA polymerase, and 19.5 µL nuclease free water. The PCR was performed with an initial denaturation at 98°C for 30 sec, then 25 cycles of denaturation at 98°C for 10 sec, annealing at 62°C for 10 sec, and elongation at 72°C for 5 min. A final elongation was performed at 72°C for 5 min. DNA cleanup, concentration measurement, purity and quality were checked as described above.

#### Cloning amplicons into plasmids

The purified PCR products were cloned into pCR-XL-2-TOPO™ vectors using the TOPO™ XL-2 Complete PCR Cloning Kit (Invitrogen). After incubation, 96 colonies were picked from the agar plate, and each transferred to its own well in a 96-well growth plate containing 1 mL of LB broth (Sigma-Aldrich) with 50 µg/mL kanamycin. The cultures were incubated at 37°C at 300 rpm for 23 h. Plasmids were purified from each culture using the QIAprep Spin Miniprep Kit (Qiagen). The concentration and quality of the DNA were checked as previously described.

#### Library preparation for Nanopore sequencing of plasmids

Library preparation was performed using the PCR barcoding (96) amplicons kit (SQK-LSK110) (Oxford Nanopore Technologies) in accordance with the manufacturer’s protocol. The first PCR amplified the target rRNA operon inserts and added an overhang used in the second PCR (barcoding PCR). The first reaction contained 20 ng of purified plasmids, 10 µL of 5X SuperFi buffer, 1 µL of 10 mM dNTP mix, 2.5 µL of 10 µM Nanopore-T3 primer, 2.5 µL of 10 µM of Nanopore-T7 primer, 0.5 µL of 2 U/µL Platinum SuperFi DNA polymerase (2 U/µL), and 19.5 µL nuclease free water. The reaction was initialised with denaturing at 98°C for 30 sec, then 10 cycles of denaturation at 98°C for 30 sec, annealing at 50°C for 30 sec, elongation at 72°C for 5 min and a final elongation at 72°C for 10 min. The PCR products were purified as previously described. The concentration and quality were checked as described above. The second PCR was performed according to supplied protocol using 0.5 nM PCR product from the first PCR reaction, 15 PCR cycles, and an extension time of 5 min. Purification, concentration and quality was checked as described above. All 96 barcoded amplicons were diluted to equimolar concentrations and pooled. The concentration and quality of the pool was checked as described above.

#### Nanopore sequencing and bioinformatics

The pooled library was sequenced using a MinION R9.4.1 flow cell on a GridION using MinKNOW v.21.11.7. The raw reads were basecalled using Guppy v.5.1.13 in super-accurate mode (dna_r9.4.1_450bps_sup.cfg). For each barcode 4000 raw reads were randomly extracted using Seqtk v.1.3 using the sample -s100 options and used for generating consensus sequences and polishing them. Canu v.2.0 (Koren et al., 2017) was used to generate consensus sequences from the raw reads using the -np genomeSize=5000 and -nanopore options. One round of polishing was made using Medaka v.1.7.2 using --model r941_min_sup_g507 for medaka consensus. After creating polished consensus sequences for each barcoded sequence the sequences were oriented using Usearch v.11.0.667 (Edgar, 2010) using a modified PR^2^ database (Jamy et al., 2022). Two rounds of orientation were made. First orientation was made using the PR^2^ database. Then a second orientation was made for the unoriented sequences using the oriented reads from the first orientation as a database. To trim the oriented consensus sequences, an alignment was made using Muscle v.3.8.31 (Edgar, 2004). The alignment was loaded into CLC Genomics Workbench v.20.0 (QIAGEN) and trimmed manually according to the primer positions in the alignment. The 18S rRNA gene was used for taxonomic classification using the PR^2^ database. Barrnap v.0.9 was used to find 18S genes using *--reject 0.4 --kingdom EUK* and Mothur v.1.41 (Schloss et al., 2009) was used to extract the sequences containing 18S genes using *get.seqs*. The 18S gene was extracted from the operons using a custom perl script (extract_18S-28S.pl) from Jamy *et al*., (2020) and used for taxonomic classification. The taxonomy was then found using a previously established phylogenetic approach (Jamy et al., 2020). Using RAxML-ng, the best unconstrained ML tree was found from 20 trees. Taxonomy from the tree was obtained using two strategies: (1) taxonomic annotation was propagated from the closest related reference to each query, and (2) all queries were removed from the tree before placing them back one at a time using EPA-ng whereafter taxonomic assignments and confidences were computed. Lastly, a consensus taxonomy for all queries was found by combining the two strategies (Table S1).

#### Plasmid linearization

Linearization was performed with suitable restriction enzymes found using CLC Genomics Workbench v.20.0 based on the criteria that they should not cleave within the insert sequences. The digestion reactions contained 77 ng plasmid DNA, 1 µL of FastDigest restriction enzyme (Fermentas), 2 µL 10X FastDigest buffer and nuclease free water to 20 µL. The following restriction enzymes and digestion conditions were used. NcoI: 10 min at 37°C and 15 min at 65°C, NotI: 30 min at 37°C and 5 min at 80°C, PstI: 5 min at 37°C and placed directly on ice, PvulI: 10 min at 37°C and placed directly on ice and ScaI: 5 min at 37°C and 10 min at 65 °C.

Before pooling the linearized plasmids into a mock community, quality checks were made to avoid potential contaminants obtained through the experimental procedure such as mixing of clones when transferring colonies from solid to liquid medium as well as by-products from the linearization. To test for mixed colonies, we mapped the oriented raw reads to the polished consensus sequences. Samples with two or more peaks were excluded as this could indicate contamination. We also checked our linearization process for unspecific digestion. This was done using gel electrophoresis, and any sequences that had more than one product were excluded from the mock community. All samples that only contained a single band were cleaned-up and pooled in equimolar concentrations to create a synthetic eukaryotic mock community. This resulted in a mock community with 41 different sequences from 46 clones, many being highly different but also some being highly similar (Figure S2). A few of the pooled mock rRNA operons were found to be 100% identical (Clone23/45, Clone40/55, and Clone30/76/77/78).

### PCR amplification of rRNA operons for benchmarking

Two separate PCR amplifications of rRNA operons were made for both the eukaryotic mock community and Askov soil sample. One for PacBio sequencing where conventional primers were used and one for nanopore sequencing using primers with UMIs (Karst et al., 2021).

#### Amplification for PacBio sequencing

PCR reactions were made with 10 µL of 5X SuperFi Buffer, 1 µL of 10 mM dNTP mix, 2.5 µL of 10 µM 3NDf primer, 2.5 µL of 10 µM of 21R primer, 10 µL SuperFi GC enhancer, 0.5 µL of 2 U/µL Platinum SuperFi DNA polymerase, either 100 ng Askov soil DNA or 0.1-0.2 ng of mock sample DNA, and nuclease free water up to 50 µL. The PCR was performed with an initial denaturation at 98°C for 30 sec, then 25 cycles of denaturation at 98°C for 10 sec, annealing at 62 for 10 sec, and elongation at 72°C for 2 min 15 sec. A final elongation was performed at 72 °C for 5 min. The PCR product was purified, and the concentration and quality were determined. The two replicates from 0.2 ng and 0.1 ng input DNA from the mock community were pooled as well as five replicates all from 100 ng input DNA from the Askov soil sample to obtain enough DNA for PacBio sequencing.

#### Amplification for nanopore sequencing

A two-step PCR approach was used to first UMI-tag rRNA operons in the samples and subsequent amplify these tagged molecules. The first PCR reactions contained 10 µL of 5X SuperFi buffer, 1 µL of 10 mM dNTP mix, 2.5 µL of 10 µM 3NDf_UMI primer, 2.5 µL of 10 µM of 21R_UMI primer, 0.5 µL of 2 U/µL Platinum SuperFi DNA polymerase, and either 0.2 ng DNA from our eukaryotic mock community or 100 ng DNA from Askov soil, and nuclease free water up to 50µL. A two cycle PCR was set up with initial denaturation at 95°C for 3 min, then 2 cycles of denaturation at 95°C for 30 sec, annealing at 62°C for 30 sec, and elongation at 72°C for 5 min. DNA clean-up was performed as previously described except the DNA was eluted in 21 µL nuclease-free water.

The second PCR reaction contained 20 µL of 5X SuperFi buffer, 2 µL of 10 mM dNTP mix, 5 µL of 10 µM lu_pcr_fw primer, 5 µL of lu_pcr_rv primer, 20 µL of template DNA from first PCR, 1 µL of 2 U/µL Platinum SuperFi DNA polymerase and 47 µL of nuclease free water. The PCR was carried out with an initial denaturation at 95°C for 3 min, then 25 cycles of denaturation at 95°C for 15 sec, annealing at 62°C for 15 sec, elongation at 72°C for 2 min 15 sec, and a final elongation at 72°C for 5 min. DNA clean-up was performed as previously described except the DNA was eluted in 22 µL nuclease-free water. The DNA concentration and quality was determined as previously described. To obtain enough material for Nanopore library preparation, the PCR product from the Askov soil sample was amplified a second time using 8 additional cycles using the same conditions as the first amplification.

After amplification of the UMI tagged molecules some DNA from each sample was used to create clonal libraries (UMI_clonal). Dilutions were made to obtain 100,000 molecules/µL as input for the amplification of the UMI_clonal library. The PCR reaction contained 20 µL of 5X SuperFi buffer, 2 µL of 10 mM dNTP mix, 5 µL of 10 µM lu_pcr_fw primer, 5 µL of 10 µM of lu_pcr_rv primer, 1 µL of template DNA (100,000 copies/µL), 1 µL of 2 U/µL Platinum SuperFi DNA polymerase, and 66 µL of nuclease-free water. The program initiated with denaturation at 95°C for 3 min, then 25 cycles of denaturation at 95°C for 30 sec, annealing at 62°C for 15 sec, elongation at 72°C for 2 min 15 sec, and a final elongation at 72°C for 5 min. DNA clean-up was performed as previously described except the DNA was eluted in 22 µL nuclease-free water. The DNA concentration and quality were determined as previously described.

The amplification of DNA for nanopore sequencing resulted in two different library types; one with amplified UMI tagged molecules (UMI library) and one with an additional amplification constructing a clonal library (UMI_clonal library).

### PacBio sequencing of rRNA operons

The two samples amplified with conventional primers were sent for PacBio library preparation and sequencing at Uppsala Genome Center, Science for Life Laboratory. Library preparation was performed using SMRTbell Express Template Prep Kit 2.0 and the samples were barcoded using Barcoded Overhang Adapter kit 8A/B. The pooled sample contained 85% mock community and 15% soil sample. One SMRT cell on a PacBio Sequel II instrument was used for sequencing the two samples using the Sequel II sequencing kit 2.0.

Circular consensus sequences (CCS) were constructed using SMRT Link v.11.0.0.146107. The CCS were trimmed, oriented and filtered using DADA2 v.1.24.0 (Callahan et al., 2016) in R v.4.2.1 (R Core Team, 2021). The reads were quality filtered as described in Jamy *et al*. 2020. Briefly, primers were removed using *removePrimers* with default settings, which also orients the reads. The reads were filtered using *filterAndTrim* with *minLen=3000, maxLen=6000* and *rm.phix=FALSE.* We observed that some reads were not properly oriented based on the above procedure and an additional orientation was therefore made with the Vsearch v.2.22.1 (Rognes et al., 2016) *-orient* command using first the modified PR^2^ database (Jamy et al., 2022) and secondly oriented reads from the first orientation as reference database. After the second orientation 100% of the reads were correctly oriented.

### Nanopore sequencing of rRNA operons

Libraries for nanopore sequencing was prepared using the samples amplified with primers containing UMIs and the Amplicons by Ligation kit (SQK-LSK110, Oxford Nanopore Technologies) according to manufacturer’s protocol. All samples were sequences individually on MinION R9.4.1 flow cells on a GridION. However, different versions of the MinKNOW software and Guppy were used as the samples were not sequenced at the same time. MinKNOW v.21.11.7 was used for the Askov soil samples (UMI and UMI_clonal) and basecalling was done with Guppy v.5.1.13 in super-accurate mode (dna_r9.4.1_450bps_sup.cfg). MinKNOW v.21.06.13 was used for the mock community UMI library and basecalling was done with Guppy v5.0.16 in super-accurate mode (dna_r9.4.1_450bps_sup.cfg). MinKNOW v.22.07.4 was used or the mock community UMI_clonal library and basecalling done with Guppy v6.3.1 in super-accurate mode (dna_r9.4.1_450bps_sup.cfg). The raw nanopore reads were processed and polished using the longread_umi pipeline (Karst et al., 2021). Primer positions were first found using the *primer_position* command with default settings. The output files “reads_t1_umi_endpos.txt” and “reads_t2_umi_endpos.txt” were used to determine the -s and -e parameters for generating UMI consensus sequences, respectively. The parameters were calculated as the mean + standard deviation + 5 of the identified positions. UMI consensus sequences were obtained using the following command *nanopore_pipeline -v 7 -q r941_min_super_g507 -m 3000 -M 6000 -c 3 -p 2 -T 2 -U “3;2;6;0.3”* and the -s and -e parameters from above. The UMI consensus sequences were oriented with Vsearch using the same approach as for the PacBio data.

### Resolving ASVs based on PacBio CCS and Nanopore UMI reads

ASV was resolved using either Unoise3 (Edgar, 2016b), Uchime3 (Edgar, 2016a) or DADA2 (Callahan et al., 2016). To ensure fair comparisons between the tested sequencing technologies, rarefraction was applied to normalize to the lowest number of consensus sequences present in the datasets using Seqtk v.1.3 (https://github.com/lh3/seqtk) with the *subseq* command.

For Unoise3 sequences was processed with Usearch v.11.0.667 (Edgar, 2010). The CCS or UMI consensus sequences were first dereplicating using the -*fastx_uniques* command with -*sizeout* and - *relabel uniq* and then denoised to resolved ASVs and remove chimera using the -*unoise3* command with *-minsize* ranging from 2 to 8.

For Uchime3 sequences were processed using Vsearch v.2.22.1 (Rognes et al., 2016). The CCS or UMI consensus sequences was first dereplicated with the --*derep_fulllength* command using --*sizeout* and --*relabel uniq* and then denoised to resolve ASVs using the -*cluster_unoise* command with --*minsize* ranging from 2 to 8, chimera was finally detected and removed using the --*uchime3_denovo* command. For DADA2 sequences were processed using DADA2 v.1.24.0 (Callahan et al., 2016) in R v.4.2.1 (R Core Team, 2021). The CCS or UMI consensus sequences was first dereplicated with the *derepFastq* command with *qualityType = “FastqQuality”*. Error models were found using *learnErrors* with *errorEstimationFunction=PacBioErrfun, randomize=TRUE, BAND_SIZE=32, qualityType=“FastqQuality”*. ASVs were resolved using *dada* with *BAND_SIZE=32.* De novo chimera detection was performed using *isBimeraDenovo* with *minFoldParentOverAbundance* ranging from 2 to 8. Because DADA2 requires fastq files as input for the *derepFastq* command. The UMI consensus fasta sequences were converted to artificial fastq files with a uniform Qscore of 40 using a custom perl command (Qscore40_script.pl) before the dereplication.

To determine the number of CCS or UMI consensus sequences and ASVs with exact matches in the mock community reference database, mappings were made using Usearch -*usearch_global* with *-id 0 - maxrejects 0 -maxaccepts 0 -strand plus* and *-top_hit_only*. To determine the number of CCS and UMI consensus sequences associated with each clone in our mock community ASV tables were made using Usearch -*otutab* command with the dereplicated sequences as input. To evaluate the de novo chimera detection, reference-based chimera detection was performed on the de novo chimera-filtered ASV using Vsearch with the *--uchime_ref* command and our mock community sequences as reference database.

### Taxonomic classification of rRNA operons from the Askov soil sample

We combined ASVs from the three Askov datasets (PacBio, and Nanopore UMI/UMI_clonal). ASVs were inferred using Unoise3 with *-minsize 2* for the Nanopore UMI data and *-minsize 3* for the Nanopore UMI_clonal and PacBio CCS data. Unique ASVs were found using Usearch with the *-fastx_uniques* command. Taxonomy was assigned using Blastn against a modified PR^2^ database (Jamy et al., 2022). Only the top hit for each query were used. Prokaryotes were removed using Mothur v.1.48.0 (Schloss et al., 2009) using the *remove.seqs* command. The sequences were aligned using MAFFT v.7.490 (Katoh et al., 2002) with *--retree 2 --maxiterate 1000 --reorder --adjustdirection* settings. The alignment was trimmed using TrimAl v1.4.1 with *-gt 0.01 -st 0.001* settings. To increase ease of visualisation, we taxonomically constrained eukaryotic groups that are well-supported in phylogenomic studies, following Jamy et al. 2022. Briefly, groups at the Supergroup, Division, and Subdivision levels in the PR^2^ database were constrained to be monophyletic, except for Excavates. A maximum likelihood phylogeny was then inferred with RAxML v.8.2.12 (Stamatakis, 2014) using the faster GTRCAT model.

### Data analysis and visualisation

The data was analysed and visualised in R v.4.1.0 (R Core Team, 2021) using R studio IDE (RStudio Team, 2020) with tidyverse v.1.3.2 (Wickham et al., 2019), ggplot2 v.3.4.0 (Wickham, 2016), data.table v.1.14.6 (Dowle & Srinivasan, 2023), dplyr v.1.0.10 (Wickham et al., 2022) and ggtree v.3.2.1 (Yu et al., 2017). Figures were assembled and polished using Adobe Illustrator v.26.3.1.

## Results

### Establishment of a synthetic eukaryotic mock community

Using mock communities has become a well-established method for benchmarking molecular methods and bioinformatic workflows. There are commercially available mock communities containing both bacterial and fungal species such as ZymoBIOMICS HMW DNA standard (Zymo Research) or custom-made mock communities from DSMZ. However, there are currently no commercially available mock communities for microeukaryotes including protists. As we wanted to evaluate different sequencing strategies and bioinformatic workflows for obtaining high-quality whole rRNA operons from microeukaryotes, we needed a well-defined mock community as reference for our analyses. Therefore, we developed a synthetic eukaryotic mock community by cloning PCR-amplified rRNA operons from soil eukaryotes into plasmids, which were subsequently replicated in competent *E. coli* cells (Figure 1). Plasmids were isolated from 96 colonies and Nanopore sequenced. Despite the relatively high error rate associated with Nanopore raw reads, we were able to generate accurate reference sequences by calculating and polishing the consensus sequences from 4000 raw reads for each clone.

**Figure 1:**
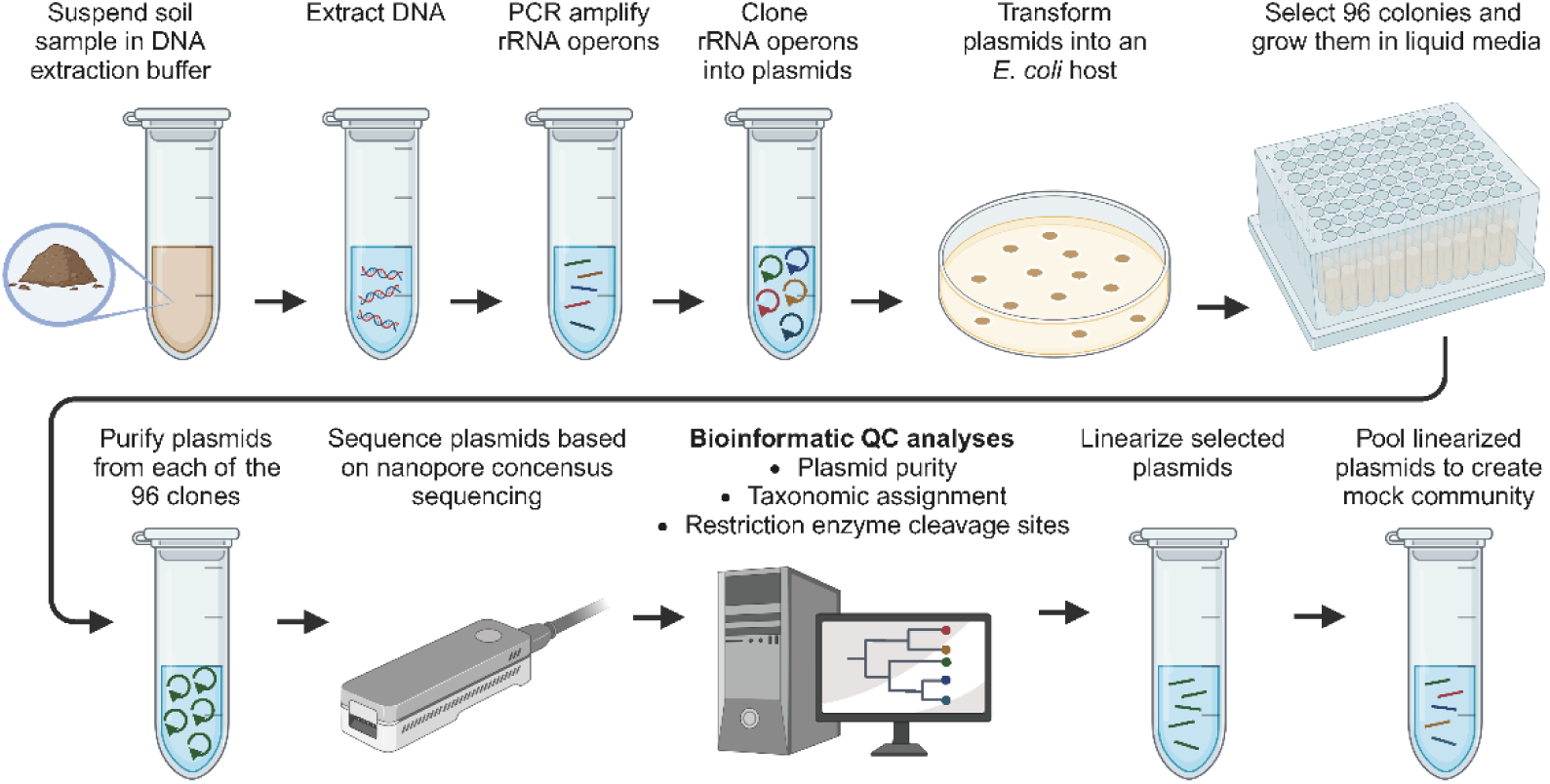
Construction of the synthetic eukaryotic mock community. DNA was extracted from soil, and eukaryotic rRNA operons were amplified using PCR. These amplified rRNA operons were then cloned into plasmids, which were subsequently transformed into *E. coli* cells. Afterwards, 96 individual colonies were transferred to liquid medium in a 96-well plate. Plasmids were purified from each of the 96 cultures and individually sequenced using Nanopore sequencing with barcoding. Consensus sequences for the rRNA operon were derived for each clone, which were then taxonomically classified. This taxonomic classification ensured a high taxonomic diversity in the final mock community. Following classification, each of the 96 plasmid samples was linearized using restriction enzymes. Those that met quality standards were pooled to create the final mock community.

We investigated the taxonomic diversity of the clone library to ensure that the mock community resembled real environmental diversity with both diverse and highly similar sequences (Figure S2). High diversity was obtained combining species from many of the lineages commonly observed in soil (Geisen et al., 2018). Fungi was the most abundant group mainly represented by Dikarya (12 taxa) but also contains Chytrids (two taxa) and Mucoromycetes (two taxa). Cercozoans and Gyristans were also abundant, represented by eight and six taxa, respectively. It is not surprising that the Dikarya has the highest abundance as this group is highly abundant in many soils (Aslani et al., 2022; Delgado-Baquerizo et al., 2018; Shen et al., 2014).

### Evaluation of long-read sequencing strategies and bioinformatic pipelines

In this study, we aimed to compare different methods for obtaining high-quality microeukaryotic rRNA operons using PacBio CCS and Nanopore UMI consensus reads (Cornaby et al., 2022; Jamy et al., 2022; Karst et al., 2021; Sereika et al., 2022). Two distinct types of Nanopore UMI libraries were created (Figure 2): Nanopore UMI, in which UMI-tagged sequences underwent a single round of amplification, and Nanopore UMI_clonal, in which UMI-tagged sequences were amplified twice. The rationale for creating the Nanopore UMI_clonal library was to optimize the sequencing capacity. By ensuring an ideal number of UMI-tagged molecules are available, we can maintain the integrity of the sequencing process. Both excessively low and high counts of tagged molecules can drastically affect the yield of UMI consensus sequences. However, it’s important to note that the second amplification step in the Nanopore UMI_clonal process might result in the introduction of chimeras, which ideally shouldn’t appear in the UMI consensus sequences. In comparison, while the PacBio CCS sequencing also runs the risk of chimera inclusion, it doesn’t benefit from the extra chimera filtering offered by UMI-tagging.

**Figure 2:**
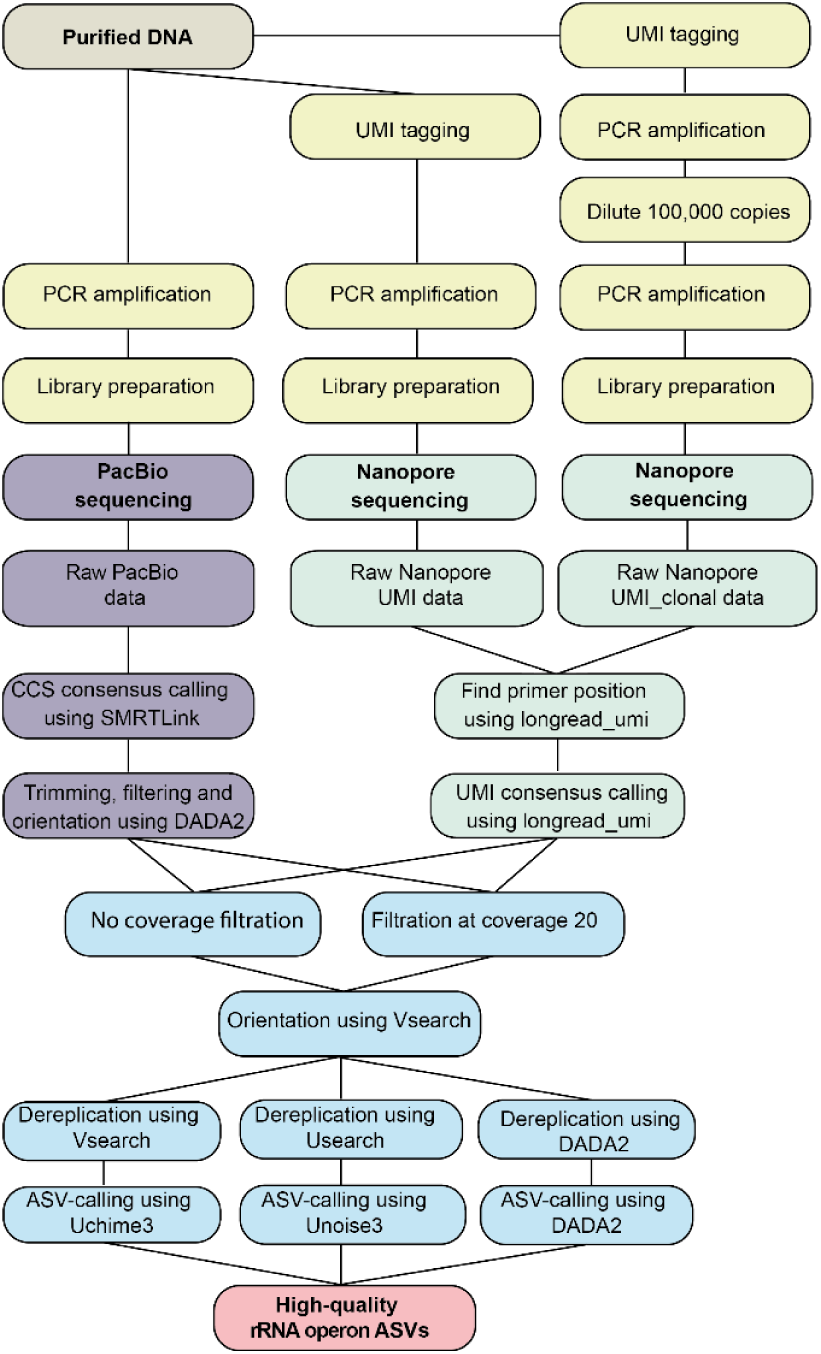
Library preparation and bioinformatic pipeline for PacBio and Nanopore data. The preparation of sequencing libraries for the different sequencing approaches are shown in yellow. The unique processing for PacBio reads is depicted in purple. CCS consensus sequences were generated from raw PacBio data using SMRT Link. Following this step, the sequences were processed by trimming the primers, performing quality filtering, and orientation using DADA2. The unique processing for Nanopore reads is depicted in green. The raw Nanopore data was processed using the longread_umi pipeline. The primer positions were first identified using the *primer_position* command, after which consensus sequences were generated using the *nanopore_pipeline* command. The resulting consensus sequences for both PacBio CCS and Nanopore UMI data were then subjected to the bioinformatic pipelines, shown in blue. First, either no coverage filtering or removal of data with coverage of less 20 was made. The two datasets from each sequencing method were then oriented by Vsearch, followed by dereplication with Vsearch, Usearch or DADA2. ASV-calling was performed using Uchime3, Unoise3 or DADA2.

For the mock community, we obtained 1,250,905 PacBio CCS reads after quality filtering, and 169,632 and 41,016 Nanopore consensus sequences for the UMI and UMI_clonal dataset, respectively. To achieve a fair comparison in the following analyses, we rarefied all datasets to 41,016 sequences. Among all the sequencing methods employed, PacBio CCS provided the best coverage of the mock community with raw sequences for 40 out of 42 taxa. In contrast, the Nanopore methods only provided coverage for 21 and 32 sequences for the UMI and UMI_clonal datasets, respectively (Figure 3a).

**Figure 3:**
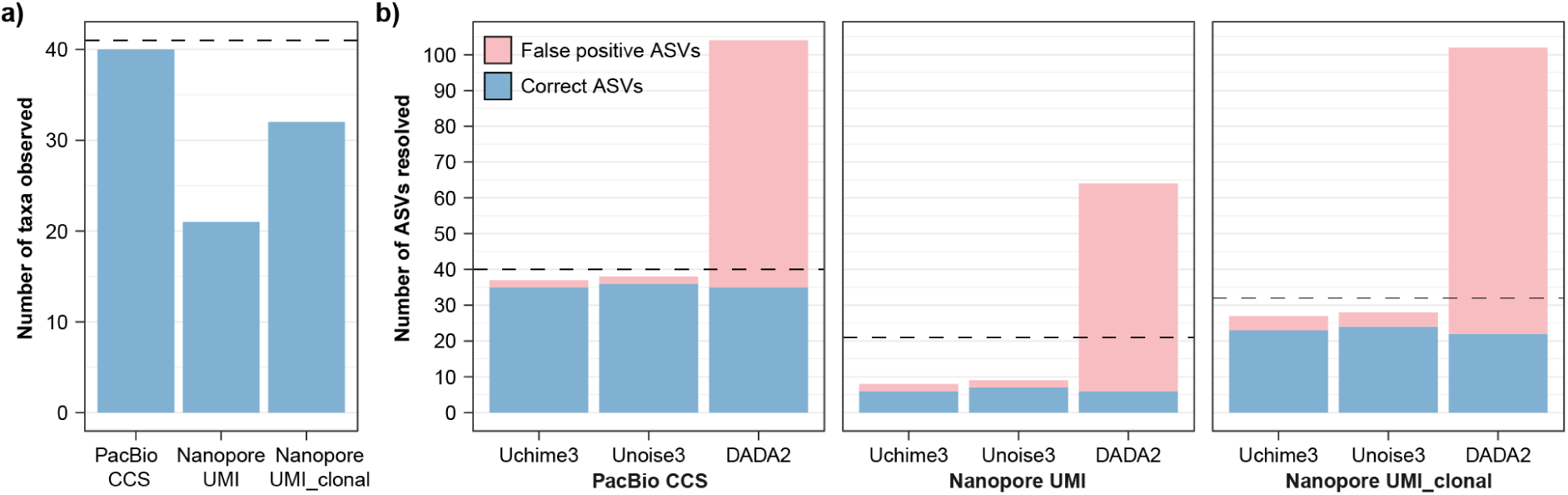
Identification of ASVs in the mock community. a) Number of mock taxa represented in the raw reads for each sequencing method. The dotted line indicates the number of unique sequences within the mock community. b) Number of ASVs found by each ASV-calling software for each of the sequencing methods. The dotted lines indicate the number of mock taxa observed in the raw reads. The following settings were used for ASV-calling: PacBio CCS (minParentAbundance 3, no coverage filtering), Nanopore UMI (minsize 2, no coverage filtering), and Nanopore UMI_clonal (minsize 3, no coverage filtering).

#### Obtaining error-free ASVs for rRNA operons

To determine the best bioinformatic pipeline for each data type, we analyzed the performance of three tools commonly used for resolving ASVs in microbiome studies: Uchime3, Unoise3 and DADA2 (Figure 3b and Table S2-S4). As error-free sequences are required for resolving ASVs, we examined the percentage of CCS and UMI consensus sequences with exact matches to the mock references as a function of the number of raw reads used to generate the consensus (Figure S3). We found that accuracy of the consensus sequences was affected by the number of CCS rounds for PacBio and the number of UMI-binned raw reads for Nanopore. The percentage of perfect sequences reached a plateau around a coverage of 20 both for the PacBio and Nanopore data (Figure S3). To investigate the potential benefits of removing low coverage data, we compared ASV-calling from unfiltered data (where no coverage filtration was done) and data filtered based on this coverage (Table S2-S4). The coverage filtration removed 64%, 69% and 57% of the PacBio CCS, Nanopore UMI and Nanopore UMI_clonal reads, respectively. Interestingly, we found that coverage filtration resulted in a reduced number of accurate ASVs compared to unfiltered data, which likely reflects the lower number of sequences available for ASV-calling after the filtration.

Another crucial factor to consider when identifying ASVs is the required abundance of a unique sequence to define an ASV. In both Uchime3 and Unoise3, this parameter is termed “minsize”, with a default value of 8. In DADA2, it’s referred to as “minParentAbundance”, which is also set at a default of 8. Decreasing this parameter can potentially increase the number of ASVs; however, it may also increase the number sequences with errors. We, therefore, examined which settings would yield the highest number of correct ASVs for the mock sample while minimising false positive ASVs, including chimeras (Table S2-S4).

The highest number of correct ASVs from the mock community was found using the PacBio data where 36 out of 41 unique sequences were identified using Unoise3 (Figure 4B). As expected, more ASVs were found with decreasing minsize for Uchime3 and Unoise3. However, we observed no effect of decreasing minParentAbundance for DADA2 (Table S2). The data, when filtered based on a coverage of 20, appeared to result in an increased number of false positive ASVs for the lower minsize values (2 and 3) with both Uchime3 and Unoise3

**Figure 4:**
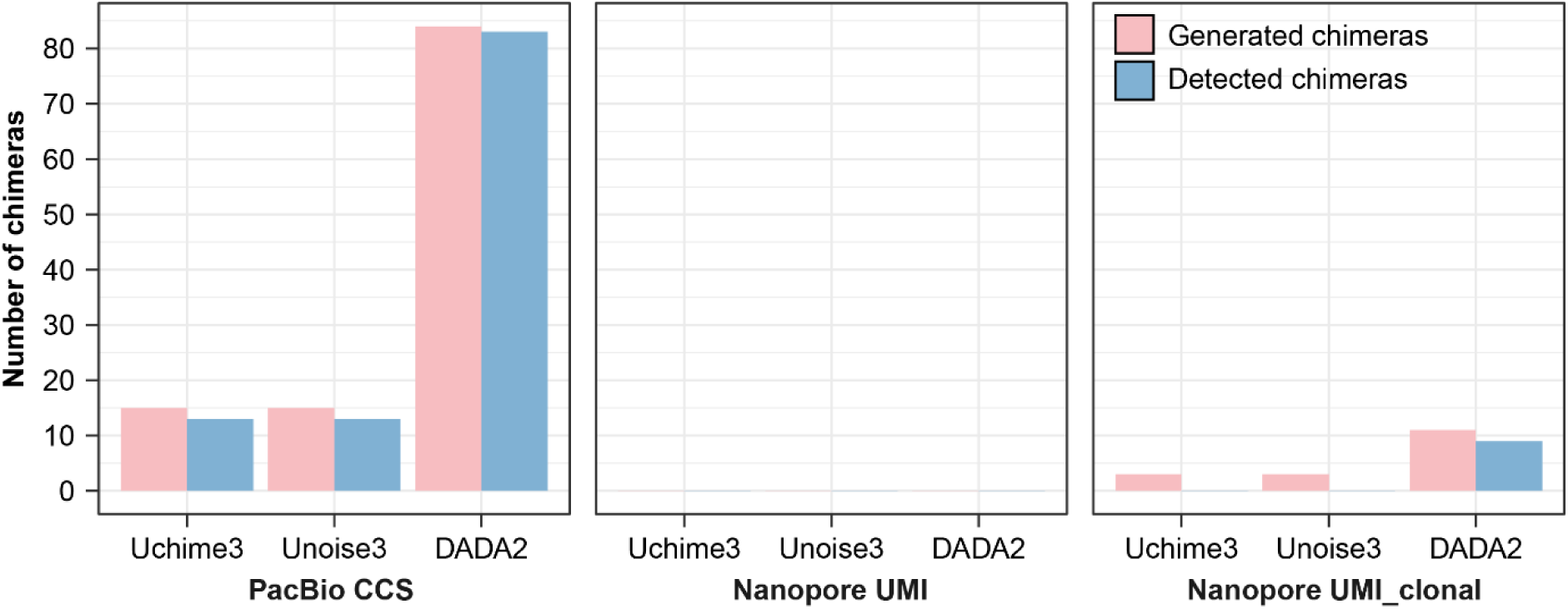
Chimera detection in the mock community. The number of chimeras found by the different chimera detection tools for each dataset. Number of chimeras are only shown for the following ASV-calling settings: PacBio CCS (minParentAbundance 3, no coverage filtering), Nanopore UMI (minsize 2, no coverage filtering), and Nanopore UMI_clonal (minsize 3, no coverage filtering).

The number of ASVs was surprisingly low for the Nanopore UMI data, with only a maximum of seven correct ASVs being identified out of 41 using Unoise3 with minsize 2 (Figure 4B). We suspect that this resulted from inefficient UMI-tagging due to the long primers used. This is supported by the observation that UMI-tagged PCR libraries frequently failed. The coverage-filtered data produced more ASVs with sequencing errors at minsize 2-4 (Table S3).

The number of correct ASVs was substantially higher for the Nanopore UMI_clonal data, with 24 correct ASVs identified out of 41 using Unoise3 with minsize 2 and 3 (Figure 4B). The coverage-filtered dataset again reduced the number of error-free ASVs, indicating that strict filtering can result in a loss of diversity (Table S4).

In all three datasets, many erroneous ASVs were observed after ASV-calling and chimera detection using DADA2, highlighting challenges in ASV-calling. While long-read sequencing methods are known to exhibit higher error rates than Illumina, most of these errors are random and can be rectified using CCS and UMIs. However, long-read sequencing methods also introduce systematic errors, one notable instance being errors due to homopolymers. The majority of noisy ASVs from DADA2 matched many of the same taxa in the mock community, differing by only a single mismatch. This suggests that the primary issue was point errors rather than homopolymers causing difficulties for DADA2.

For all three datasets, Unoise3 was able to correctly identify an ASV (clone32) that was not detected by Uchime3 and DADA2. Since this sequence differs from clone83 by only a single nucleotide, it’s likely that clone32 is predicted as a sequence error due to the settings in Uchime3 and DADA2. Given that Uchime3 generally performed very similarly to Unoise3, it represents a notable open-access alternative to the commercial Unoise3 software.

None of the methods were able to identify all 41 distinct taxa in the mock sample. To determine whether the absence of certain taxa was due to PCR amplification bias, we mapped the CCS and UMI consensus sequences to the mock reference sequences to check for missing reads corresponding to these taxa in the raw data (Figure S3). Indeed, the mock community taxa not detected at minsize 2 in the UMI dataset, or at minsize 3 in the UMI-clonal and PacBio datasets, seemed to lack adequate raw consensus reads for ASV-calling. This discrepancy in the UMI datasets could be attributed to insufficient UMI-tagging, as highlighted by the pronounced uneven coverage across various mock references.

#### Presence of remaining chimeras

To detect chimeras, we employed a two-step process. First, we used de novo chimera detection (integrated in Unoise3 and standalone in Uchime3). Secondly, we performed reference-based chimera detection on the de novo chimera-filtered ASVs using our mock reference sequences to identify any remaining chimeras. Reference-based chimera detection is highly effective when a comprehensive database that encompasses all taxa in the sample, such as in mock communities, is available. However, for environmental samples, this often isn’t the case, and de novo chimera detection becomes the primary approach.

For the PacBio data, chimeras were observed across all minsizes in varying numbers, ranging from 1 to 73. However, no chimeras were found at minsize 8 for both Unoise3 and Uchime3 when utilizing de novo chimera detection (Table S2). While the presence of chimeras in this data type is anticipated, detected chimeras are not concerning since they are filtered out. Yet, reference-based chimera detection indicated that a few undetected chimeras were present across all minsizes. As the minsize value decreased, this method revealed a rise in the number of undetected chimeras. While DADA2 identified many ASVs and likewise detected numerous chimeras, the count remained consistent regardless of decreasing minParentenAbundance. When the data was filtered based on a coverage of 20, there was an evident increase in undetected chimeras for the lower minsize values (2 and 3) with Uchime3 and Unoise3, as evidenced by the heightened chimera count in the reference-based detection for these minsizes. Utilizing minsize 7 and 8 resulted in neither chimeras nor noisy ASVs, but it also brought about a reduction in the number of detected error-free ASVs.

As anticipated, no chimeras were identified in the Nanopore UMI data by any software, whether through de novo or reference-based ASV-calling. This validates that all chimeras are removed during data processing. In light of the chimera-free nature of this data, employing minsize 2 for ASV-calling is feasible. While this data is devoid of chimeras, some might question the need for ASV-calling. However, it’s essential to acknowledge that many sequences may still contain random sequencing errors that could impact subsequent analyses. The majority of these errors are efficiently mitigated during ASV-calling.

A significant difference between the UMI_clonal and the UMI datasets was the chimera detection resulting from the second amplification of the UMI-tagged molecules. De novo chimera detection only identified chimeras at minsize 2 for Uchime3 and Unoise3, but chimeras were detected across all minParentAbundance levels for DADA2. Moreover, reference-based chimera detection highlighted a chimera that had gone undetected by de novo chimera detection. Overall, the reference-based chimera detection for the three tools revealed that ASV-calling algorithms aren’t flawless. Even in denoised data, a few unnoticed chimeras might persist unless UMIs are applied directly.

### Recommendations for obtaining high-quality rRNA operons

For the PacBio CCS data, we advocate for the use of either Uchime3 or Unoise3 with minsize 3 for ASV-calling. Given that the sequencing process and the bioinformatic pipeline for the Nanopore UMI data effectively eliminate all chimeras, it’s feasible to use Uchime3 or Unoise3 with minsize of 2. However, for Nanopore UMI_clonal data, we recommend a minsize of 3 for either Uchime3 or Unoise3, given the potential for chimera formation during the second amplification. The choice of minsize represents a balance between sequence precision and sequencing depth. Although chimeras are observed in reference-based ASV-calling for both PacBio and Nanopore UMI_clonal data at minsize 3, the count remains minimal. We anticipate a more significant loss of diversity in complex samples, compared to the mock community, should one choose a higher minsize for ASV-calling. There’s no need for CCS/UMI coverage filtering across any of the data types.

### High-quality long-read data enables the creation of rRNA operons from soil eukaryotes

To determine whether the findings from the mock community are consistent with more complex samples, we analyzed an agricultural soil sample from Askov, Denmark. From this sample, we obtained 335,947 PacBio CCS reads, 31,693 Nanopore UMI and 16,257 Nanopore UMI_clonal consensus sequences. To ensure a balanced comparison of ASV-calling and chimera filtering, all datasets were rarefied to 16,257 sequences. ASV-calling was done using the recommended settings for each data type found above.

For the PacBio data, 297 ASVs were identified, which was the lowest number compared to the other datasets (Table S5). In contrast to the mock sample, no chimeras were detected at minsize 3 in the Askov soil sample. However, undetected chimeras might still be present. The Nanopore UMI data yielded 1,022 ASVs. Despite the low number of ASVs detected in the mock community, this dataset produced the most ASVs for the soil sample (Table S5). It appears that the stochastic errors in UMI-tagging were not as pronounced for this specific sample. The ASV-calling of the Nanopore UMI_clonal data resulted in 525 ASVs. We anticipated a greater number of ASVs in the UMI_clonal sample because a second amplification increased the quantity of each uniquely tagged molecule, which could, in theory, enhance consensus calling.

### Eukaryotic diversity in an agricultural soil

To explore the microeukaryotic diversity within the Askov soil sample, we combined ASVs from the three types of sequencing data and obtained 5,123 unique near full-length rRNA operons. These operons contain the 18S, 5.8S, and 28S rRNA genes as well as the ITS1 and ITS2 regions. The rRNA operons from the Askov soil sample included representatives from nearly all major eukaryotic lineages (Figure 5). As expected, Fungi and Cercozoans were particularly diverse, followed by Ciliates, Apicomplexans, Gyristans, Chlorophytes, and Amoebozans (Geisen et al., 2014, 2018; Pellegrino et al., 2021). Predominantly marine groups such as Haptophytes, Radiolarians, and Foraminiferans were not represented.

**Figure 5:**
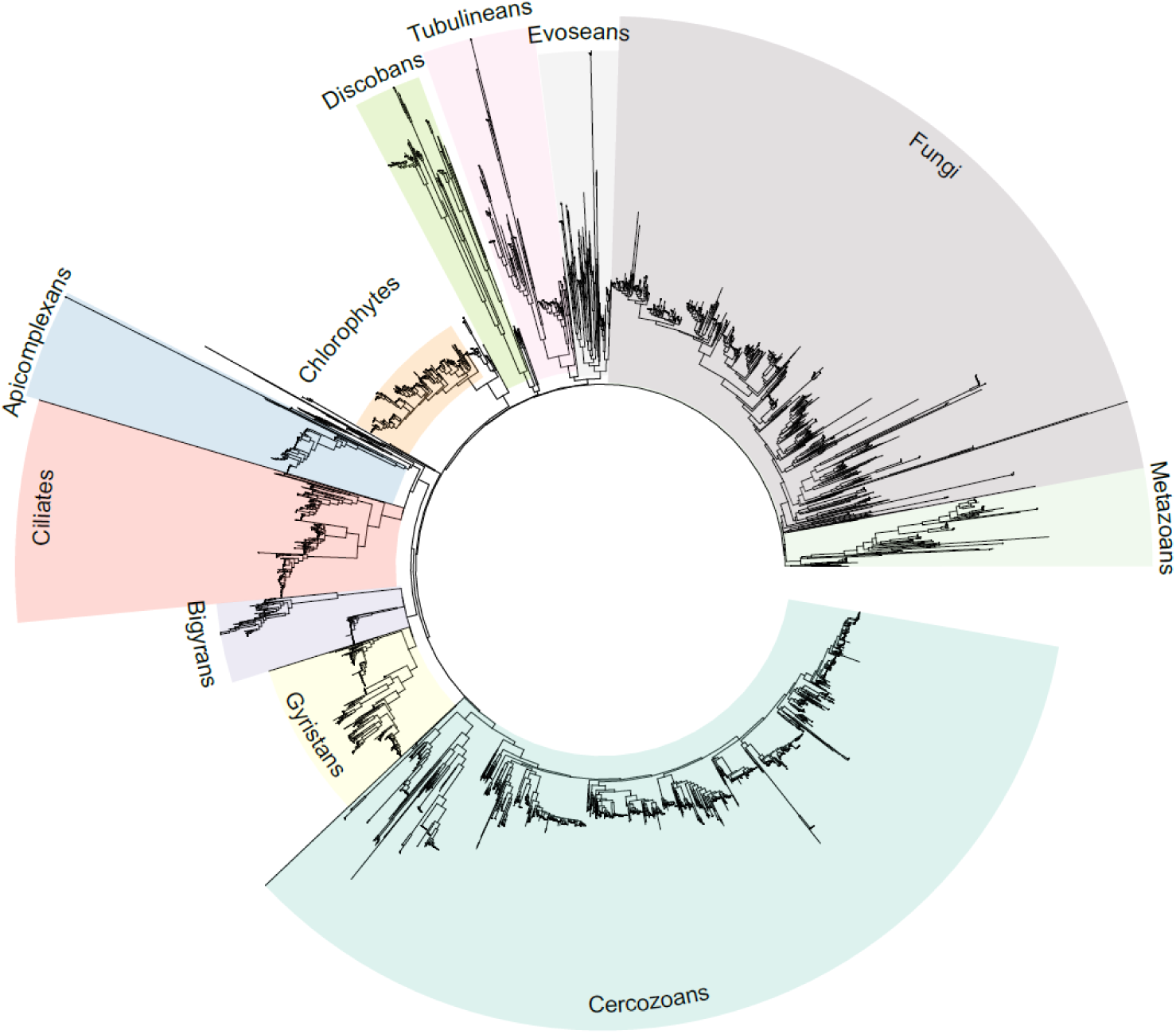
Eukaryotic taxonomic diversity within Askov soil. Constrained phylogenetic tree showing the different taxonomic groups within the Askov soil.

Our findings align with previous studies on eukaryotic diversity in soil. However, our study is the first to analyze ASV-resolved rRNA operons from microeukaryotes. This distinction is significant because the additional inclusion of ITS regions can provide greater taxonomic resolution for certain taxa.

## Discussion

In this study, we developed a method to construct a synthetic eukaryotic mock community, which was then used to provide recommendations for how to achieve highly accurate whole rRNA operons from microeukaryotes based on PacBio CCS or Nanopore UMI consensus reads. The approach was applied to a complex agricultural sample to identify 5,123 unique ASV-resolved rRNA operons providing a first glimpse into the hidden diversity of microeukaryotes within our soils.

### Challenges and considerations with Nanopore UMI sequencing

To increase the accuracy of Nanopore sequencing, we used UMIs to reduce sequencing errors. However, incorporating UMIs isn’t without its challenges. Notable among these are the stochastic biases introduced by UMI-tagging and the difficulties posed by PCR with lengthy UMI primers, leading to inconsistent results. Molecules not tagged during the initial PCR phase evade detection in subsequent analyses. It is also important to adjust the number of unique UMI-tagged molecules in the PCR amplification. Too few UMIs will result in low taxa diversity, whereas too many can lead to insufficient coverage for consensus calling. Even with adjustments, as seen in the Nanopore UMI_clonal approach, targeting the desired number of molecules can still be challenging. Thus, the use of UMIs is a compromise between obtaining chimera-free, high-quality data and potentially missing taxa. Therefore, while UMIs have been useful for generating high-quality data, the advantages in accuracy might no longer exceed the disadvantages regarding inconsistency as other sequencing methods improve.

The recent development of the R10 chemistry from Nanopore has shown promising results in enhancing nanopore sequencing accuracy. In particular, the R10 chemistry should determine homopolymer lengths more accurately. However, the current error rate still prevents the use of ASV-calling on these raw reads.

### Comparative analysis of Nanopore MinION sequencing and PacBio sequel II

To compare the sequencing methods, we evaluated various parameters ranging from sequencing cost to data processing time and data quality for both Nanopore MinION sequencing and PacBio Sequel II (Table 2). In terms of library preparation time, Nanopore library preparation using UMIs takes approximately one day, while PacBio library preparation is quicker, requiring about half a day. Both Nanopore and PacBio sequencing demand high DNA quality. However, Nanopore sequencing has stricter requirements, as contaminants from DNA extraction can damage the nanopores, thereby reducing sequencing capacity. Moreover, we’ve observed that the crucial UMI tagging reaction is significantly impacted, even by a low number of impurities.

**Table 2:** Comparison of PacBio CCS and Nanopore UMI consensus techniques for high-throughput sequencing of environmental eukaryotic rRNA operons.

|  | PacBio CCS | Nanopore UMI | Nanopore UMI_clonal |
| --- | --- | --- | --- |
| Library prep time | 0.5 day | 1 day | 1 day |
| DNA quality requirement | High | Very high | Very high |
| Raw read processing | 0-1 day | > 1 week per MinION run* | > 1 week per MinION run* |
| Reproducibility | High | Low | Low |
| Remaining sequencing errors | Very few | Very few | Very few |
| Remaining chimeras | Very few | None | Very few |
| UMI/CCS reads | 4,000,000 | up to 100,000<sup>†</sup> | up to 100,000<sup>‡</sup> |
| Estimated price raw reads | 1100€ (Sequel II) | 600€ (MinION) | 600€ (MinION) |

\* Processing time scale non-linearly with the amount of data, which prevents upscaling
<sup>†</sup> Difficult to exploit the full sequencing capacity. Number of UMIs can vary.
<sup>‡</sup> Generally more UMIs compared to Nanopore UMI

One of the limitations of these long-read sequencing technologies is the estimation of homopolymer length. It’s anticipated that both technologies will exhibit residual sequencing errors, with most non-random errors being homopolymers. This issue is particularly pronounced for nanopore data.

A clear advantage of the UMI nanopore sequencing approach is the exclusion of chimeras in the processed data. This advantage only applies to the Nanopore UMI data, not the Nanopore UMI_clonal data, since the second amplification here can result in chimera formation (Figure 4). A few of these might remain undetected after chimera detection, as observed with the PacBio data.

The cost of operating a single MinION flow cell (R9 or R10) was approximately 600€ per sample, including consumables. For sequencing the Askov and mock samples using Nanopore technology, we utilized two MinION flow cells, dedicating one to each of our Nanopore UMI libraries. The Nanopore UMI data yielded 169,632 consensus sequences for the mock community, whereas the Nanopore UMI_clonal data produced 41,016 consensus sequences. This underscores the potential variability in output for UMI datasets. It’s notable that processing raw nanopore reads demands a significant amount of computing time (>1 week on a high-performance computer) when employing the longread_umi pipeline, during which numerous temporary files are generated and processes run.

The cost of a flow cell and the library preparation kit for the new PromethION stands at approximately 1000€. Each flow cell can produce around 150 Gb of data, which is five times more than what one might expect from a high-performing MinION flow cell. However, leveraging this enhanced sequencing capacity is challenging because bioinformatic processing doesn’t scale linearly with the data volume.

We utilized PacBio Sequel II for CCS data generation, which cost approximately 1100€ per sample and yielded around 2.1 million CCS reads, compared to an expected 4 million. The recent introduction of PacBio Revio has significantly enhanced sequencing output and accuracy, allowing three times more data to be acquired from one flow cell at the same cost as the Sequel II (PacBio, 2023). Since both the Sequel IIe and Revio deliver processed CCS directly, bioinformatics processing is substantially reduced. This advantage, combined with a much lower cost per consensus sequence, positions the PacBio platform as the current method of choice.

### Towards highly accurate whole rRNA operons and improved classification methods

Accurate classification at higher taxonomic levels necessitates high-quality reference sequences in the databases. Currently, these databases rely on portions of the operon, such as the 18S rRNA, ITS, or 28S rRNA genes. By producing highly accurate whole rRNA operons, we aim to establish new standards for the taxonomic classification of microeukaryotes. However, comprehensive databases for whole rRNA operons are not yet available. To address this gap, more eukaryotic genomes, inclusive of the entire rRNA operons, are needed to secure full-length rRNA operons with genome classification. Initiatives are already underway for animals, plants, and fungi (Lewin et al., 2018). An alternative method for classifying whole rRNA operons involves the use of AutoTax. This software was developed to create ecosystem-specific 16S rRNA gene reference databases, using the SILVA taxonomy as a backbone. Additionally, it provides de novo placeholder names for lineages without an official taxonomy, ensuring a comprehensive seven-level taxonomy for all input sequences (Dueholm et al., 2020). However, when working with eukaryotes, it is crucial to contemplate potential adjustments to the software, such as opting for the PR2 database over SILVA. Moreover, the taxonomic classification threshold currently employed for prokaryotes might not directly apply to eukaryotes and warrants further exploration. Also, the use of seven taxonomic levels may not convey the same significance for eukaryotes as for prokaryotes, so this should be carefully assessed or possibly expanded.

In conclusion, the development of a pipeline for generating highly accurate whole rRNA operons from eukaryotes using advanced sequencing technologies and bioinformatic tools is critical for accurate microbial community identification. Continued research efforts to improve sequencing accuracy and data output, will be important for the development of a comprehensive eukaryotic database.

### Future studies

To deepen our understanding and refine the accuracy of our results, several additional steps can be considered. First, comparing the long-read sequencing data with short-read data from the same sample would be beneficial. Such a comparative analysis would enable us to evaluate the advantages and disadvantages of each sequencing technology, pinpointing potential discrepancies or biases between them.

Given the observed differences in UMI performance between low and high diversity mock communities, it would be compelling to study mock communities with a range of diversity levels. Such an investigation would clarify the effect of diversity on UMI efficacy and provide insights into the reliability and consistency of UMI-based tagging across various ecological contexts.

Additionally, a comparative analysis between the 18S gene sequences or short read amplicons and the entire rRNA operon sequences would be interesting. By examining the differences and similarities between these regions, we can assess the potential benefits of using full operon sequences for taxonomic classification compared to focusing solely on the 18S gene. This analysis would contribute to our understanding of the information content and discriminatory power of different regions within the rRNA operon.

### Final remarks and recommendations

Based on the findings from this study, the UMI-based approach may appear to be a promising method for studying more complex samples. However, it is important to consider the challenges associated with the amplification using UMI primers, especially given the length of these primers and stringent DNA quality demands. In contrast, the PacBio method might be preferable since it demands less labor, and showcases better robustness and reproducibility, without the stochastic bias that UMI-tagging introduces. Overall, the choice of sequencing technology and settings is a compromise between sequencing depth and potential undetected chimeras. Therefore, it is essential to consider the specific research question and the potential biases introduced by the sequencing and processing settings. The results of this analysis suggest that UMI sequencing may be advantageous in reducing the detection of chimeras, while PacBio sequencing may be more effective in detecting diversity. With the rapid advancements in sequencing technologies, it is becoming increasingly feasible to sequence the majority of all ecosystems in the near future.

## Acknowledgements

This study was funded by Independent Research Fund Denmark (Grant no. 9041-00236B).

## Data Accessibility and Benefit-Sharing

## Data Accessibility Statement

All sequencing data have been submitted to the Sequence Read Archive under the project ID PRJNA1019781. All R scripts and raw data used for data analysis and visualisation are available at GitHub: https://github.com/msdueholm/Publications/Overgaard2023.

## Benefit-Sharing

Benefits Generated: Benefits from this research accrue from the sharing of our data and results on obtaining high-quality whole rRNA operons.

## Author Contributions

CKO prepared the sequencing of whole rRNA operons as well as experimental preparation of the eukaryotic mock community and nanopore sequencing. CKO and MKDD processed the sequencing data. CKO and MJ did the bioinformatics analyses. CKO, MJ, and MKDD wrote the manuscript and CKO and MKDD designed the study. SR, FB, and MKDD supervised the project. All authors approved the final manuscript.

## Supplementary Figures

**Figure S1:**
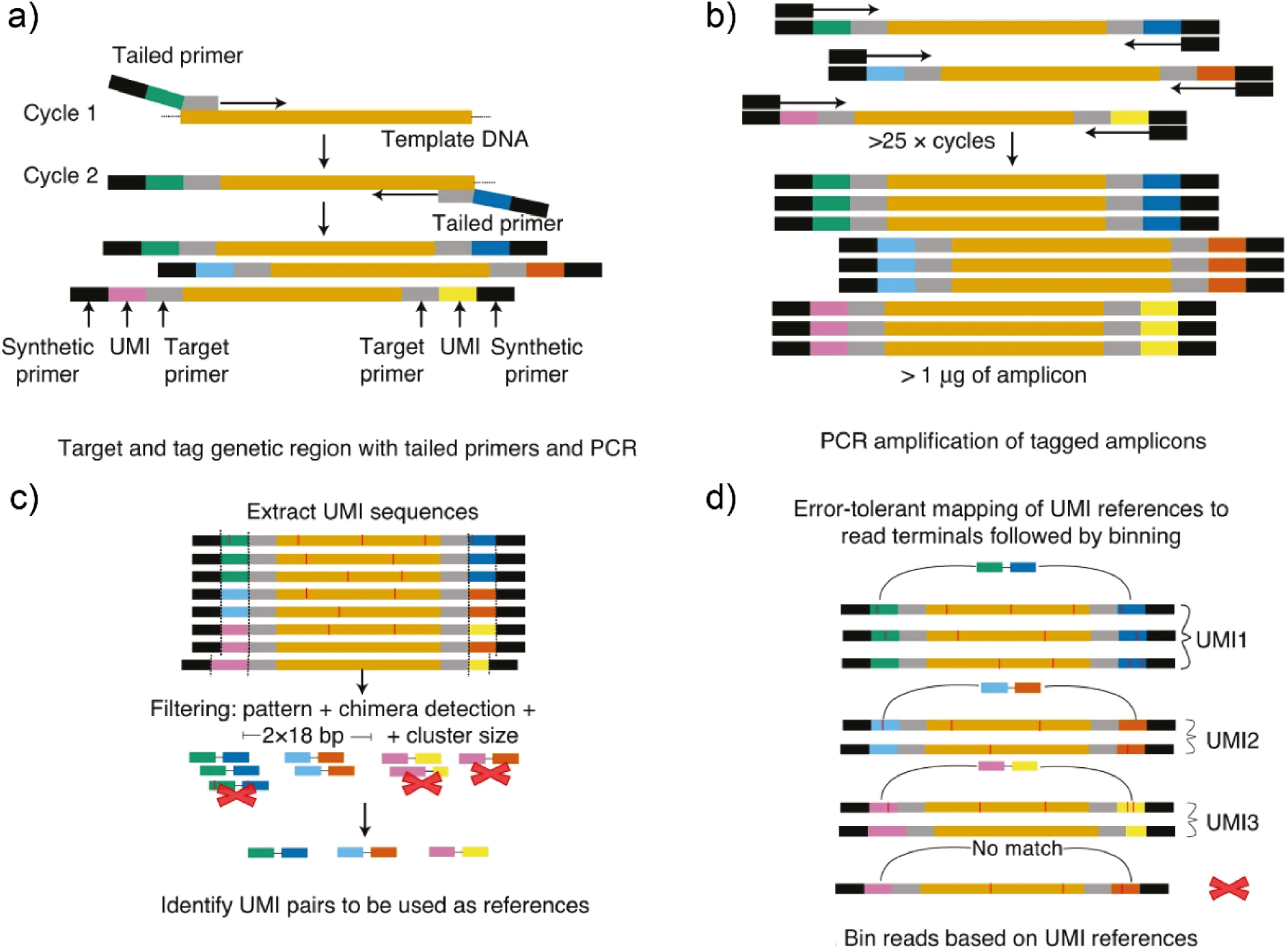
Tagging approach using unique molecular identifiers (UMIs). a) two-cycle PCR tagging reaction is used to tag target molecules with UMIs in each end of the molecules. b) Amplification of UMI-tagged molecules is used to create clonal copies of each UMI-tagged molecule. c) Chimeras are removed based on incorrect UMI pairs. d) The reads are binned according to their UMI pairs and high-quality consensus sequences are found for each bin. Revised from (Karst et al., 2021).

**Figure S2:**
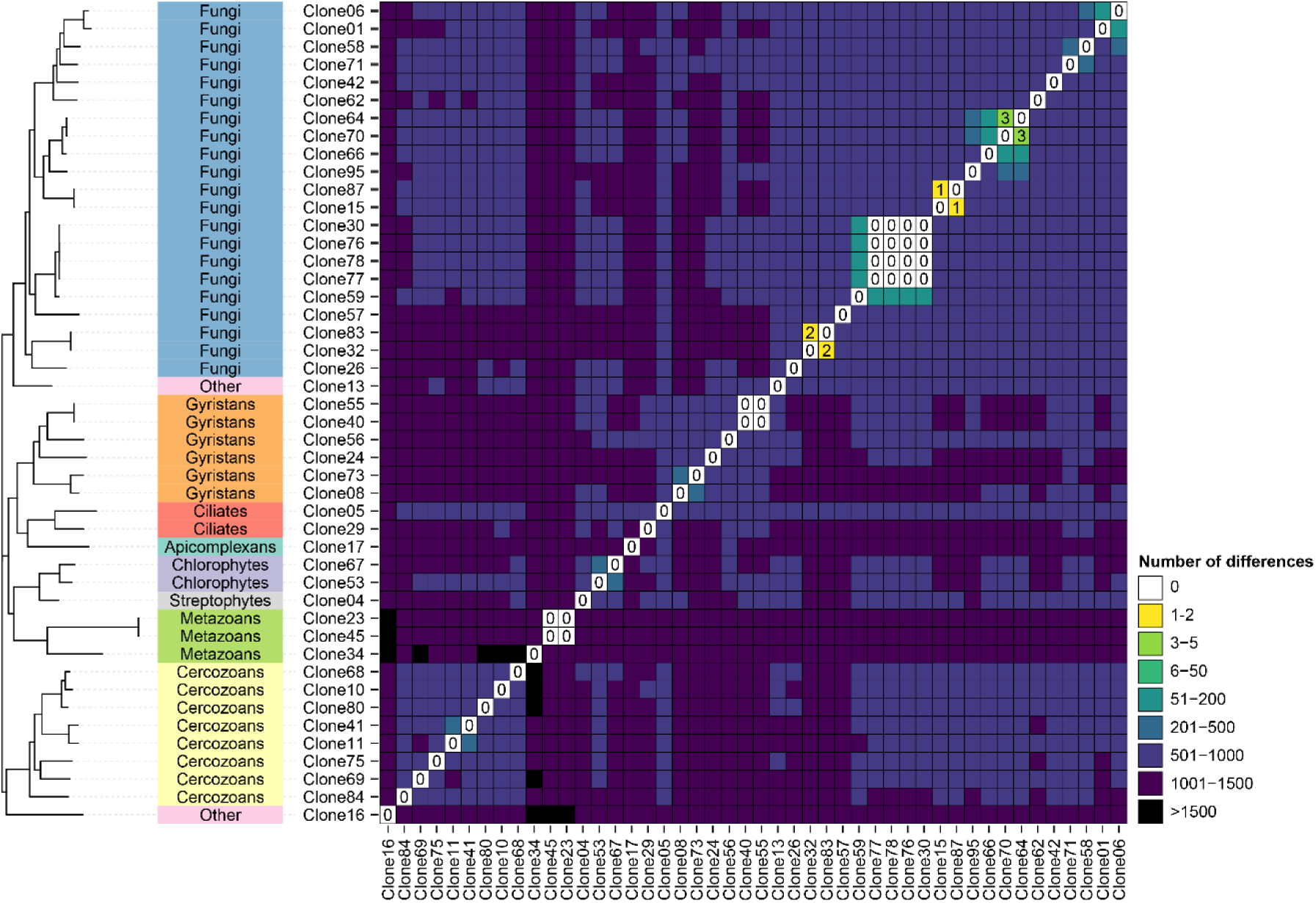
Phylogenetic analysis of rRNA operon sequences in the mock community. The names in the colored bars represent the class rank of each sequence. Colors in the heatmap indicate the number of mismatches or gaps between the 46 references in the mock community. Differences count both mismatches and gaps between the sequences. Differences below 10 are provided as text in the heatmap.

**Figure S3:**
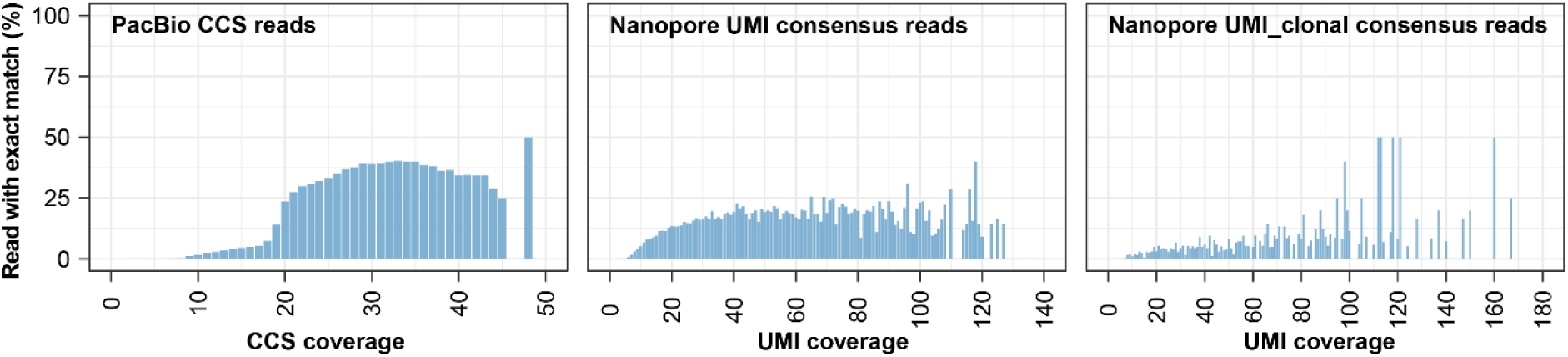
Percentage of perfect consensus sequences for each UMI/CCS coverage. Filtered and oriented PacBio CCS and Nanopore UMI consensus sequences were mapped to the mock reference sequences to find the percentage of exact hits as a function of CCS or UMI coverage.

**Figure S4:**
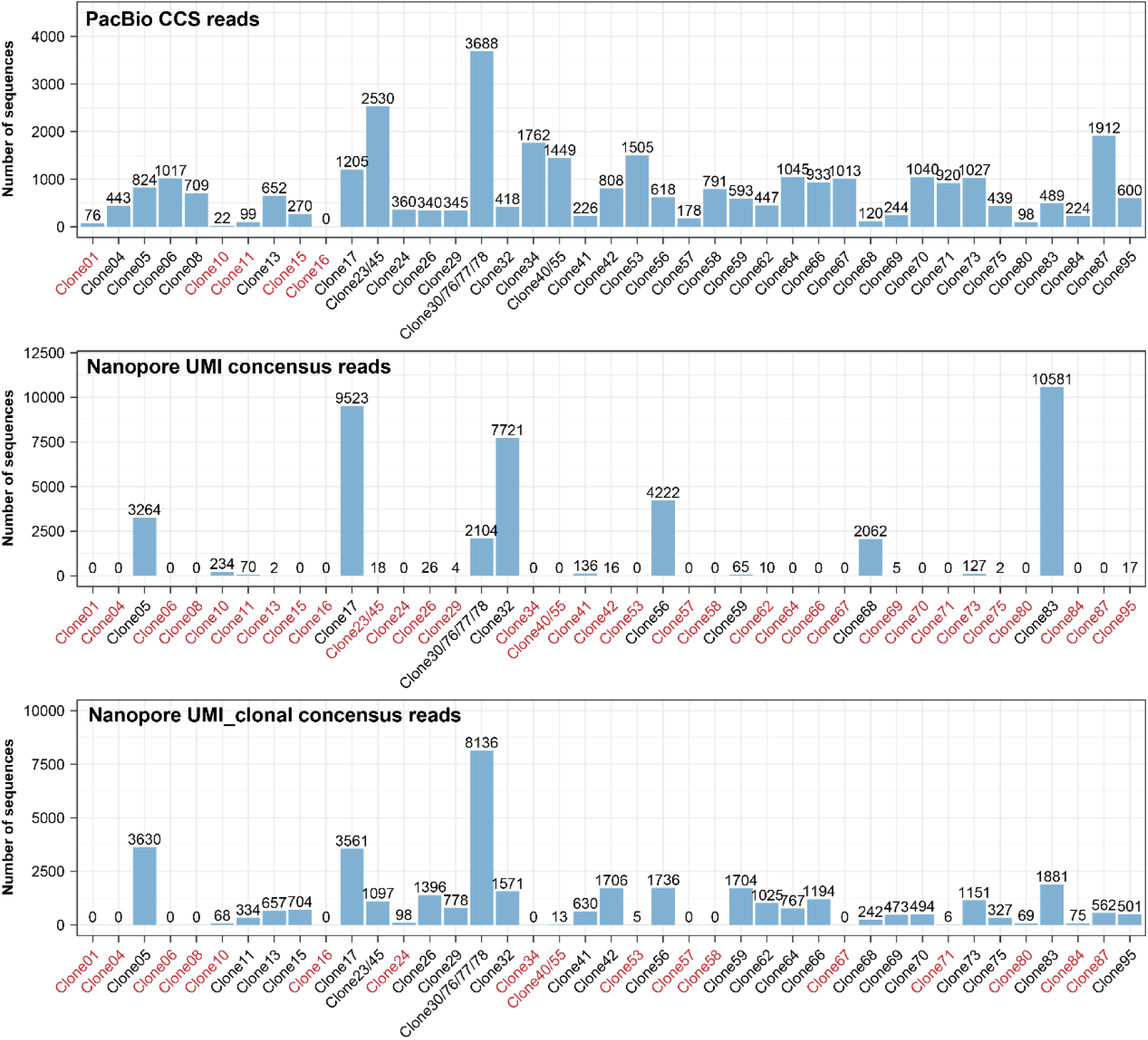
Number of sequences mapping to each of the mock reference sequences. Filtered and oriented PacBio CCS or Nanopore UMI consensus reads were mapped to the mock reference sequences for each of the data sets to investigate if CCS or UMI consensus sequences were present for taxa that were not observed after ASV-calling (marked in red).

## Tables

**Table S1:**
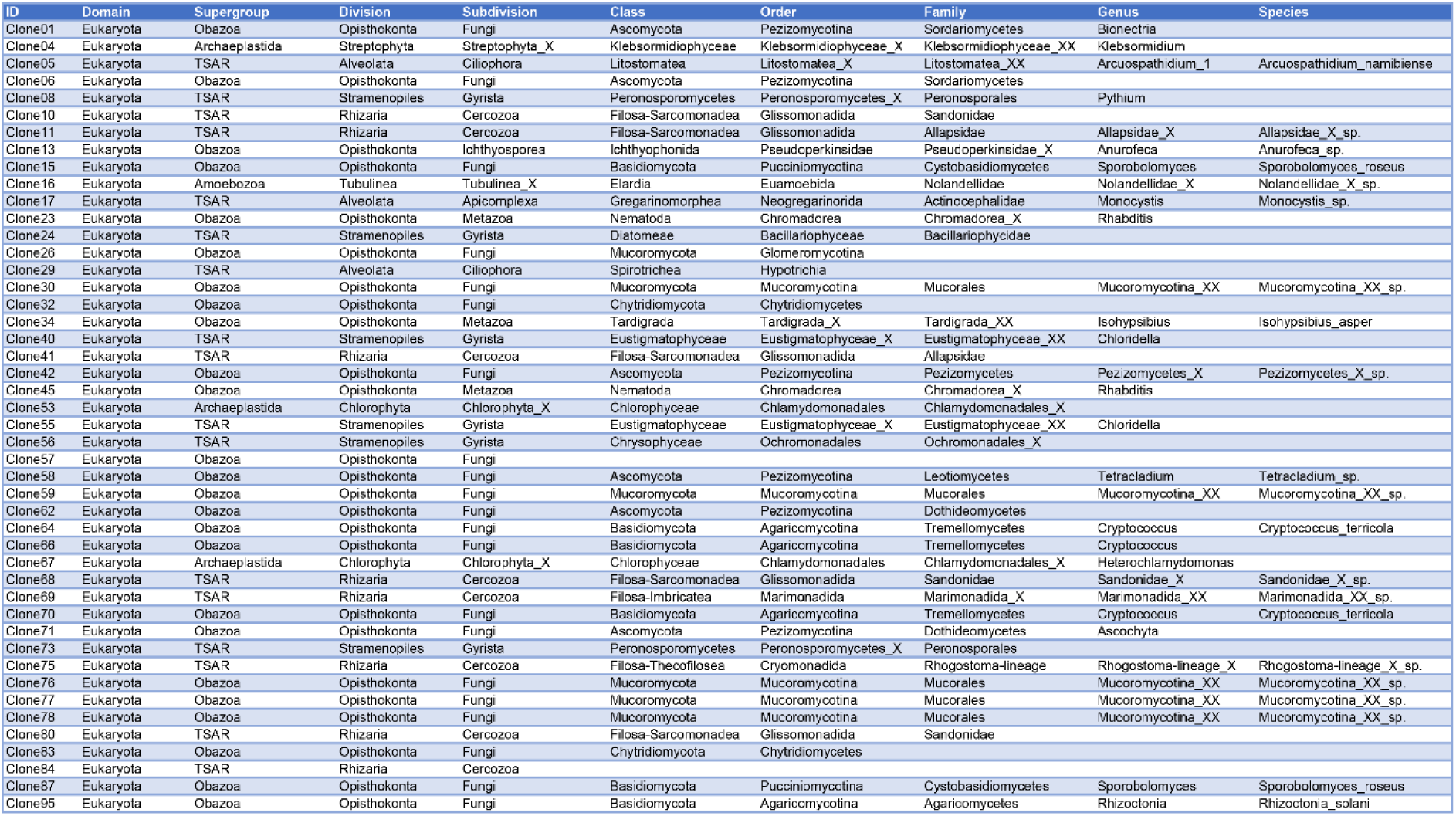
Taxonomy for the eukaryotic mock sequences obtained based on the PR^2^ database.

**Table S2:** ASV-calling and chimera detection of PacBio CCS data for the eukaryotic mock community. Three different tools were used for ASV-calling: Uchime3 (Edgar, 2016a), Unoise3 (Edgar, 2016b) and DADA2 (Callahan et al., 2016). For each tool different minimum abundances of the centroid/parent sequence were investigated. For DADA2 this parameter is called minParentAbundance (mPA) and for Uchime3 and Unoise3 it is called minsize. Unfiltered data has no CCS coverage filtering whereas Coverage 20 data was filtered to only include CCS reads with a coverage ≥ 20. Two rounds of ASV-calling and chimera detection were made. First one using the de novo method and then one using the reference method where the ASVs from the first round of ASV-calling were checked against the mock reference sequences. Detected ASVs is the amount of ASVs found by the software and error-free ASVs is the amount of ASVs that has a perfect match to the mock community. Marked in red are the settings we recommend for ASV-calling on PacBio CCS reads.

| PacBio CCS for Mock community |  |  |  |  |  |  |  |  |
| --- | --- | --- | --- | --- | --- | --- | --- | --- |
|  | Unfiltered |  |  |  | Coverage 20 |  |  |  |
|  | De novo chimera detection |  | Reference chimera detection |  | De novo chimera detection |  | Reference chimera detection |  |
|  | Detected/Error-free ASVs | Chimeras | Detected/Error-free ASVs | Chimeras | Detected/Error-free ASVs | Chimeras | Detected/Error-free ASVs | Chimeras |
| **Uchime3** |  |  |  |  |  |  |  |  |
| minsize2 | 38 / 35 | 72 | 35 / 35 | 3 | 39 / 35 | 44 | 35 / 35 | 4 |
| minsize3 | **37 / 35** | **13** | 35 / 35 | 2 | 38 / 35 | 6 | 35 / 35 | 3 |
| minsize4 | 37 / 35 | 2 | 35 / 35 | 2 | 36 / 35 | 2 | 35 / 35 | 1 |
| minsize5 | 37 / 35 | 1 | 35 / 35 | 2 | 36 / 35 | 1 | 35 / 35 | 1 |
| minsize6 | 36 / 35 | 1 | 35 / 35 | 1 | 36 / 35 | 0 | 35 / 35 | 1 |
| minsize7 | 34 / 33 | 1 | 33 / 33 | 1 | 33 / 33 | 0 | 33 / 33 | 0 |
| minsize8 | 34 / 33 | 0 | 33 / 33 | 1 | 33 / 33 | 0 | 33 / 33 | 0 |
| **Unoise3** |  |  |  |  |  |  |  |  |
| minsize2 | 38 / 36 | 73 | 36 / 36 | 2 | 40 / 36 | 44 | 37 / 36 | 3 |
| minsize3 | **38 / 36** | **13** | 36 / 36 | 2 | 39 / 36 | 6 | 36 / 36 | 3 |
| minsize4 | 38 / 36 | 2 | 36 / 36 | 2 | 37 / 36 | 2 | 36 / 36 | 1 |
| minsize5 | 38 / 36 | 1 | 36 / 36 | 2 | 37 / 36 | 1 | 36 / 36 | 1 |
| minsize6 | 37 / 36 | 1 | 36 / 36 | 1 | 37 / 36 | 0 | 36 / 36 | 1 |
| minsize7 | 35 / 34 | 1 | 34 / 34 | 1 | 34 / 34 | 0 | 34 / 34 | 0 |
| minsize8 | 35 / 34 | 0 | 34 / 34 | 1 | 34 / 34 | 0 | 34 / 34 | 0 |
| **DADA2** |  |  |  |  |  |  |  |  |
| mPA2 | 104 / 35 | 83 | 103 / 35 | 1 | 76 / 35 | 51 | 76 / 35 | 0 |
| mPA3 | 104 / 35 | 83 | 103 / 35 | 1 | 76 / 35 | 51 | 76 / 35 | 0 |
| mPA4 | 104 / 35 | 83 | 103 / 35 | 1 | 76 / 35 | 51 | 76 / 35 | 0 |
| mPA5 | 104 / 35 | 83 | 103 / 35 | 1 | 76 / 35 | 51 | 76 / 35 | 0 |
| mPA6 | 104 / 35 | 83 | 103 / 35 | 1 | 76 / 35 | 51 | 76 / 35 | 0 |
| mPA7 | 104 / 35 | 83 | 103 / 35 | 1 | 76 / 35 | 51 | 76 / 35 | 0 |
| mPA8 | 104 / 35 | 83 | 103 / 35 | 1 | 76 / 35 | 51 | 76 / 35 | 0 |

**Table S3:** ASV-calling and chimera detection of Nanopore UMI data for the eukaryotic mock community. Three different tools were used for ASV-calling: Uchime3 (Edgar, 2016a), Unoise3 (Edgar, 2016b) and DADA2 (Callahan et al., 2016). For each tool different minimum abundances of the centroid/parent sequence were investigated. For DADA2 this parameter is called minParentAbundance (mPA) and for Uchime3 and Unoise3 it is called minsize. Unfiltered data has no coverage filtering whereas Coverage 20 data was filtered to only include reads with a coverage ≥ 20. Two rounds of ASV-calling and chimera detection were made. First one using the de novo method and then one using the reference method where the ASVs from the first round of ASV-calling were checked against the mock reference sequences. Detected ASVs is the amount of ASVs found by the software and error-free ASVs is the amount of ASVs that has a perfect match to the mock community. Marked in red are the settings we recommend for ASV-calling for Nanopore UMI data.

| Nanopore UMI for Mock community |  |  |  |  |  |  |  |  |
| --- | --- | --- | --- | --- | --- | --- | --- | --- |
|  | Unfiltered |  |  |  | Coverage 20 |  |  |  |
|  | De novo chimera detection |  | Reference chimera detection |  | De novo chimera detection |  | Reference chimera detection |  |
|  | Detected/Error-free ASVs | Chimeras | Detected/Error-free ASVs | Chimeras | Detected/Error-free ASVs | Chimeras | Detected/Error-free ASVs | Chimeras |
| **Uchime3** |  |  |  |  |  |  |  |  |
| minsize2 | 8 / 6 | 0 | 8 / 6 | 0 | 10 / 6 | 0 | 10 / 6 | 0 |
| minsize3 | 6 / 5 | 0 | 6 / 5 | 0 | 8 / 6 | 0 | 8 / 6 | 0 |
| minsize4 | 6 / 5 | 0 | 6 / 5 | 0 | 7 / 6 | 0 | 7 / 6 | 0 |
| minsize5 | 6 / 5 | 0 | 6 / 5 | 0 | 5 / 5 | 0 | 5 / 5 | 0 |
| minsize6 | 6 / 5 | 0 | 6 / 5 | 0 | 5 / 5 | 0 | 5 / 5 | 0 |
| minsize7 | 6 / 5 | 0 | 6 / 5 | 0 | 5 / 5 | 0 | 5 / 5 | 0 |
| minsize8 | 6 / 5 | 0 | 6 / 5 | 0 | 5 / 5 | 0 | 5 / 5 | 0 |
| **Unoise3** |  |  |  |  |  |  |  |  |
| minsize2 | 9 / 7 | 0 | 9 / 7 | 0 | 11 / 7 | 0 | 11 / 7 | 0 |
| minsize3 | 7 / 6 | 0 | 7 / 6 | 0 | 9 / 7 | 0 | 9 / 7 | 0 |
| minsize4 | 7 / 6 | 0 | 7 / 6 | 0 | 8 / 7 | 0 | 8 / 7 | 0 |
| minsize5 | 7 / 6 | 0 | 7 / 6 | 0 | 6 / 6 | 0 | 6 / 6 | 0 |
| minsize6 | 7 / 6 | 0 | 7 / 6 | 0 | 6 / 6 | 0 | 6 / 6 | 0 |
| minsize7 | 7 / 6 | 0 | 7 / 6 | 0 | 6 / 6 | 0 | 6 / 6 | 0 |
| minsize8 | 7 / 6 | 0 | 7 / 6 | 0 | 6 / 6 | 0 | 6 / 6 | 0 |
| **DADA2** |  |  |  |  |  |  |  |  |
| mPA2 | 64 / 6 | 0 | 64 / 6 | 0 | 6 / 6 | 0 | 6 / 6 | 0 |
| mPA3 | 64 / 6 | 0 | 64 / 6 | 0 | 6 / 6 | 0 | 6 / 6 | 0 |
| mPA4 | 64 / 6 | 0 | 64 / 6 | 0 | 6 / 6 | 0 | 6 / 6 | 0 |
| mPA5 | 64 / 6 | 0 | 64 / 6 | 0 | 6 / 6 | 0 | 6 / 6 | 0 |
| mPA6 | 64 / 6 | 0 | 64 / 6 | 0 | 6 / 6 | 0 | 6 / 6 | 0 |
| mPA7 | 64 / 6 | 0 | 64 / 6 | 0 | 6 / 6 | 0 | 6 / 6 | 0 |
| mPA8 | 64 / 6 | 0 | 64 / 6 | 0 | 6 / 6 | 0 | 6 / 6 | 0 |

**Table S4:** ASV-calling and chimera detection of Nanopore UMI_clonal data for the eukaryotic mock community. Three different tools were used for ASV-calling: Uchime3 (Edgar, 2016a), Unoise3 (Edgar, 2016b) and DADA2 (Callahan et al., 2016). For each tool different minimum abundances of the centroid/parent sequence were investigated. For DADA2 this parameter is called minParentAbundance (mPA) and for Uchime3 and Unoise3 it is called minsize. Unfiltered data has no coverage filtering whereas Coverage 20 data was filtered to only include reads with a coverage ≥ 20. Two rounds of ASV-calling and chimera detection were made. First one using the de novo method and then one using the reference method where the ASVs from the first round of ASV-calling were checked against the mock reference sequences. Detected ASVs is the amount of ASVs found by the software and error-free ASVs is the amount of ASVs that has a perfect match to the mock community. Marked in red are the settings we recommend for ASV-calling for Nanopore UMI_clonal data.

| Nanopore UMI_clonal for Mock community |  |  |  |  |  |  |  |  |
| --- | --- | --- | --- | --- | --- | --- | --- | --- |
|  | Unfiltered |  |  |  | Coverage 20 |  |  |  |
|  | De novo chimera detection |  | Reference chimera detection |  | De novo chimera detection |  | Reference chimera detection |  |
|  | Detected/Error-free ASVs | Chimeras | Detected/Error-free ASVs | Chimeras | Detected/Error-free ASVs | Chimeras | Detected/Error-free ASVs | Chimeras |
| **Uchime3** |  |  |  |  |  |  |  |  |
| minsize2 | 40 / 23 | 8 | 35 / 23 | 5 | 22 / 16 | 4 | 20 / 16 | 2 |
| minsize3 | **27 / 23** | **0** | 24 / 23 | 3 | 15 / 14 | 0 | 14 / 14 | 1 |
| minsize4 | 21 / 21 | 0 | 21 / 21 | 0 | 12 / 12 | 0 | 12 / 12 | 0 |
| minsize5 | 19 / 19 | 0 | 19 / 19 | 0 | 12 / 12 | 0 | 12 / 12 | 0 |
| minsize6 | 17 / 17 | 0 | 17 / 17 | 0 | 11 / 11 | 0 | 11 / 11 | 0 |
| minsize7 | 17 / 17 | 0 | 17 / 17 | 0 | 10 / 10 | 0 | 10 / 10 | 0 |
| minsize8 | 16 / 16 | 0 | 16 / 16 | 0 | 9 / 9 | 0 | 9 / 9 | 0 |
| **Unoise3** |  |  |  |  |  |  |  |  |
| minsize2 | 42 / 24 | 7 | 37 / 24 | 5 | 23 / 17 | 4 | 21 / 17 | 2 |
| minsize3 | **28 / 24** | **0** | 25 / 24 | 3 | 16 / 15 | 0 | 15 / 15 | 1 |
| minsize4 | 22 / 22 | 0 | 22 / 22 | 0 | 13 / 13 | 0 | 13 / 13 | 0 |
| minsize5 | 20 / 20 | 0 | 20 / 20 | 0 | 13 / 13 | 0 | 13 / 13 | 0 |
| minsize6 | 18 / 18 | 0 | 18 / 18 | 0 | 12 / 12 | 0 | 12 / 12 | 0 |
| minsize7 | 18 / 18 | 0 | 18 / 18 | 0 | 11 / 11 | 0 | 11 / 11 | 0 |
| minsize8 | 17 / 17 | 0 | 17 / 17 | 0 | 10 / 10 | 0 | 10 / 10 | 0 |
| **DADA2** |  |  |  |  |  |  |  |  |
| mPA2 | 102 / 22 | 9 | 100 / 22 | 2 | 33 / 16 | 4 | 32 / 16 | 1 |
| mPA3 | 102 / 22 | 9 | 100 / 22 | 2 | 33 / 16 | 4 | 32 / 16 | 1 |
| mPA4 | 102 / 22 | 9 | 100 / 22 | 2 | 33 / 16 | 4 | 32 / 16 | 1 |
| mPA5 | 102 / 22 | 9 | 100 / 22 | 2 | 33 / 16 | 4 | 32 / 16 | 1 |
| mPA6 | 102 / 22 | 9 | 100 / 22 | 2 | 33 / 16 | 4 | 32 / 16 | 1 |
| mPA7 | 102 / 22 | 9 | 100 / 22 | 2 | 33 / 16 | 4 | 32 / 16 | 1 |
| mPA8 | 102 / 22 | 9 | 100 / 22 | 2 | 33 / 16 | 4 | 32 / 16 | 1 |

**Table S5:** ASV-calling and chimera detection of Nanopore UMI and CCS data for eukaryotic rRNA operons from Askov soil. Three different tools were used for ASV-calling: Uchime3 (Edgar, 2016a), Unoise3 (Edgar, 2016b) and DADA2 (Callahan et al., 2016). For each tool different minimum abundances of the centroid/parent sequence were investigated. For DADA2 this parameter is called minParentAbundance (mPA) and for Uchime3 and Unoise3 it is called minsize. Unfiltered data has no read coverage filtering whereas Coverage 20 data was filtered to only include reads with a coverage ≥ 20. Two rounds of ASV-calling and chimera detection were made: one using the de novo method and one using the reference method where the ASVs from the de novo ASV-calling were ASV-called and chimera checked against the mock reference sequences. Two rounds of ASV-calling and chimera detection were made. First one using the de novo method and then one using the reference method where the ASVs from the first round of ASV-calling were checked against the mock reference sequences. Detected ASVs is the amount of ASVs found by the software and error-free ASVs is the amount of ASVs that has a perfect match to the mock community. Marked in red are the settings we recommend for ASV-calling for the different datasets.

|  | PacBio CCS for Askov soil samples |  |  |  |  | Nanopore UMI for Askov soil samples |  |  |  |  | Nanopore UMI_clonal for Askov soil samples |  |  |  |
| --- | --- | --- | --- | --- | --- | --- | --- | --- | --- | --- | --- | --- | --- | --- |
|  | Unfiltered |  | Coverage 20 |  |  | Unfiltered |  | Coverage 20 |  |  | Unfiltered |  | Coverage 20 |  |
| Uchime3 | ASVs | Chimeras | ASVs | Chimeras |  | ASVs | Chimeras | ASVs | Chimeras |  | ASVs | Chimeras | ASVs | Chimeras |
| minsize2 | 554 | 2 | 481 | 1 |  | 1017 | 0 | 922 | 0 |  | 1015 | 0 | 953 | 0 |
| minsize3 | 296 | 0 | 255 | 0 |  | 526 | 0 | 481 | 0 |  | 520 | 0 | 483 | 0 |
| minsize4 | 208 | 0 | 183 | 0 |  | 358 | 0 | 382 | 0 |  | 355 | 0 | 325 | 0 |
| minsize5 | 157 | 0 | 138 | 0 |  | 269 | 0 | 242 | 0 |  | 281 | 0 | 255 | 0 |
| minsize6 | 130 | 0 | 113 | 0 |  | 213 | 0 | 189 | 0 |  | 219 | 0 | 198 | 0 |
| minsize7 | 104 | 0 | 92 | 0 |  | 175 | 0 | 150 | 0 |  | 173 | 0 | 161 | 0 |
| minsize8 | 88 | 0 | 80 | 0 |  | 145 | 0 | 131 | 0 |  | 154 | 0 | 141 | 0 |
| Unoise3 |  |  |  |  |  |  |  |  |  |  |  |  |  |  |
| minsize2 | 556 | 3 | 481 | 2 |  | 1022 | 0 | 926 | 0 |  | 1023 | 0 | 960 | 0 |
| minsize3 | 297 | 0 | 225 | 0 |  | 529 | 0 | 484 | 0 |  | 525 | 0 | 488 | 0 |
| minsize4 | 208 | 0 | 183 | 0 |  | 361 | 0 | 331 | 0 |  | 360 | 0 | 330 | 0 |
| minsize5 | 157 | 0 | 138 | 0 |  | 272 | 0 | 245 | 0 |  | 285 | 0 | 259 | 0 |
| minsize6 | 130 | 0 | 113 | 0 |  | 215 | 0 | 191 | 0 |  | 222 | 0 | 200 | 0 |
| minsize7 | 104 | 0 | 92 | 0 |  | 177 | 0 | 151 | 0 |  | 175 | 0 | 163 | 0 |
| minsize8 | 88 | 0 | 80 | 0 |  | 146 | 0 | 132 | 0 |  | 156 | 0 | 143 | 0 |
| DADA2 |  |  |  |  |  |  |  |  |  |  |  |  |  |  |
| mPA2 | 562 | 5 | 488 | 4 |  | 1042 | 4 | 896 | 3 |  | 907 | 3 | 838 | 3 |
| mPA3 | 562 | 5 | 488 | 4 |  | 1042 | 4 | 896 | 3 |  | 907 | 3 | 838 | 3 |
| mPA4 | 562 | 5 | 488 | 4 |  | 1042 | 4 | 896 | 3 |  | 907 | 3 | 838 | 3 |
| mPA5 | 562 | 5 | 488 | 4 |  | 1042 | 4 | 896 | 3 |  | 907 | 3 | 838 | 3 |
| mPA6 | 562 | 5 | 488 | 4 |  | 1042 | 4 | 896 | 3 |  | 907 | 3 | 838 | 3 |
| mPA7 | 562 | 5 | 488 | 4 |  | 1042 | 4 | 896 | 3 |  | 907 | 3 | 838 | 3 |
| mPA8 | 562 | 5 | 488 | 4 |  | 1042 | 4 | 896 | 3 |  | 907 | 3 | 838 | 3 |

## Notes

https://github.com/msdueholm/Publications/tree/master/Overgaard2023

https://www.ebi.ac.uk/ena/browser/view/PRJNA1019781

### Competing Interest Statement

The authors have declared no competing interest.

